# Ultra-flexible endovascular probes for brain recording through micron-scale vasculature

**DOI:** 10.1101/2023.03.20.533576

**Authors:** Anqi Zhang, Emiri T. Mandeville, Lijun Xu, Creed M. Stary, Eng H. Lo, Charles M. Lieber

## Abstract

Implantable neuroelectronic interfaces have enabled significant advances in both fundamental research and treatment of neurological diseases, yet traditional intracranial depth electrodes require invasive surgery to place and can disrupt the neural networks during implantation. To address these limitations, we have developed an ultra-small and flexible endovascular neural probe that can be implanted into small 100-micron scale blood vessels in the brains of rodents without damaging the brain or vasculature. The structure and mechanical properties of the flexible probes were designed to meet the key constraints for implantation into tortuous blood vessels inaccessible with existing techniques. *In vivo* electrophysiology recording of local field potentials and single-unit spikes has been selectively achieved in the cortex and the olfactory bulb. Histology analysis of the tissue interface showed minimal immune response and long-term stability. This platform technology can be readily extended as both research tools and medical devices for the detection and intervention of neurological diseases.

## Main

Neuroelectronic interfaces establish communication between the brain and external devices (*1-3*). Many such interfaces have been developed to gather and modulate different forms of neural information, yet there is a clear trade-off between invasiveness and precision. Specifically, non-penetrating techniques such as electroencephalography (EEG) (*4*) and electrocorticography (ECoG) (*5*) are less invasive, but lack the spatial resolution to target individual neurons and are limited to recording from the brain surface. In contrast, invasive approaches such as depth electrodes (*6, 7*) can achieve single-cell, single-spike resolution in deep brain regions, but the open-skull implantation pose considerable risks (*8*), including intracortical bleeding and infection, as well as damage to the targeted brain regions. To overcome the trade-off of invasiveness and resolution, here we report an ultra-flexible micron-scale neuroelectronic interface that utilizes a native delivery system: the brain vasculature. The metabolically active central nervous system demands a dense vascular network, so the average neuron is less than 20 μm from the nearest blood vessel (*9, 10*). This vasculature thus offers recording probes access to any brain region without damaging the recorded neuron circuits.

Endovascular recording of brain waves in millimeter-scale blood vessels has been demonstrated in previous studies (*11, 12*). In 1973, the first endovascular EEG in human (*13*) was achieved using a guided stainless-steel catheter with a 1.5 mm × 0.6 mm electrode. Through the internal carotid artery (ICA) in the neck, the catheter was advanced to the middle cerebral artery (MCA) in the brain (with a diameter of ∼ 2.9 mm (*14*)). The state-of-the-art endovascular recording electrode, stentrode™ (Synchron) was first reported in 2016 and later commercialized and approved for clinical trials, has an electrode array consisting of 8 electrodes with 0.75-mm electrode discs fabricated on a self-expanding stent (*15, 16*). The stent was inserted through the external jugular vein in the neck into superior sagittal sinus (with a diameter of ∼ 2.4 mm) in sheep, and achieved chronic recording for up to 28 weeks. However, much of the brain remains inaccessible with these electrodes, and endovascular recording has never been achieved in small rodent animal models, because the metal-based catheters and stents are stiff and bulky, and navigating them through tortuous brain vasculature with down to micron-scale vessels can result in tissue damage and inflammation. Other reports proposed the use of smaller devices to avoid tissue damage. For example, a platinum electrode on the tip of a micrometer-scale platinum wire could be loaded into the vessels by the saline flow, and was demonstrated for *ex vivo* recording inside intestinal and spinal cord vessels of frogs (*17*). Polymer-based wires with a magnetic head can also be driven with flow and magnetic steering, and demonstrated navigation inside microfluidic devices and showed endovascular insertion in *ex vivo* rabbit ears (*18*). However, *in vivo* endovascular electrophysiology has not yet been achieved with such flexible devices. To enable basic neuroscience research and clinical practice, we need to have the probes capable of *in vivo* electrophysiology applicable first to the micron-scale vasculature in small rodent animal models, and then scale to larger animals such as humans.

Here, we demonstrate ultra-flexible micro-endovascular (MEV) probes that can be precisely delivered through the blood vessels in the neck into small 100-micron scale vessels in rat brains, without damaging the brain or vasculature (Fig. 1a). Using our MEV probes, *in vivo* electrophysiology recording of local field potentials and single-unit spikes has been selectively achieved in the cortex and the olfactory bulb. The flexible probe/vessel wall/brain tissue interface exhibits minimal inflammatory response and long-term stability.

**Figure 1.**
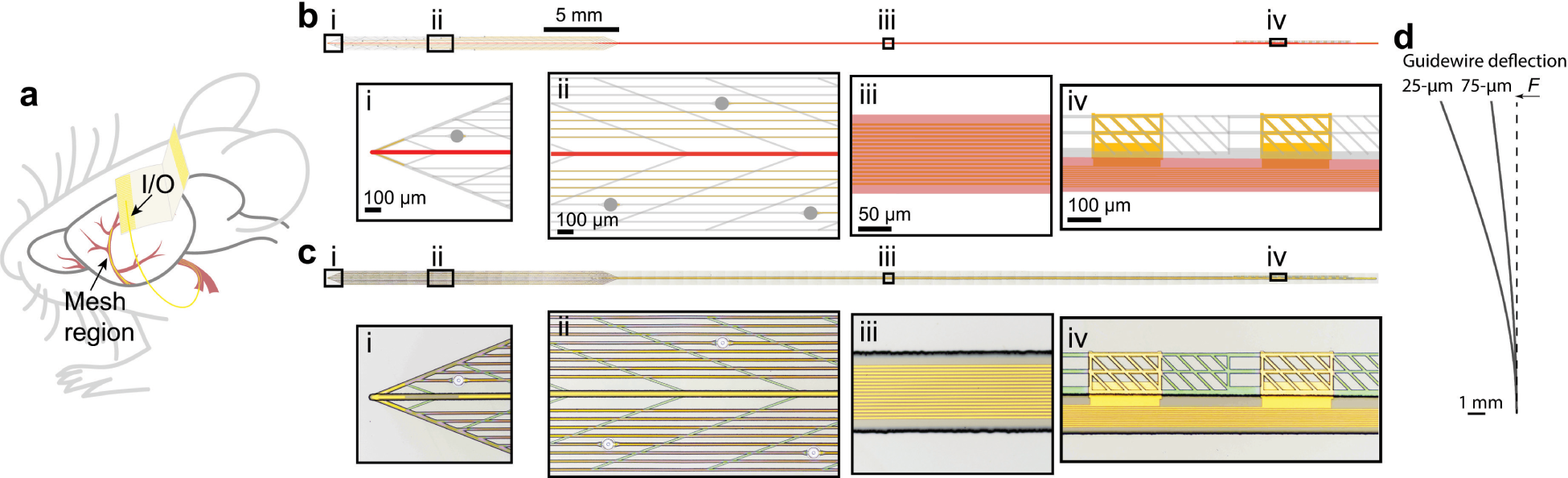
Endovascular implantation and overview of the MEV probe. **a**, Schematic showing an MEV probe (yellow) implanted into a rat brain through the blood vessels in the neck. The mesh region with electrodes is implanted deeper, while the input/output (I/O) region remains exteriorized for subsequent connection and measurement. **b**, Schematic of the MEV probe with the ultra-flexible mesh-like device region at left tapering into the stem in the middle and I/O region at right. The total length of the probe is 7 cm. The middle line in red is the thick SU-8 guidewire layer. Insets provide magnified views of (i) tip of the device region containing the 25-μm wide guidewire (red) and the ultra-flexible SU-8 mesh structured region (light gray), (ii) Pt recording electrodes (gray circles) supported on the SU-8 mesh structured region (light gray), (iii) the SU-8 stem containing independent metal interconnects for each electrode, and (iv) the I/O pads consist of gold region (yellow) connected by SU-8 mesh structured region (light gray). **c**, Tiled bright-field optical microscopy images of a representative MEV probe with a 25-μm wide guidewire in the device region. Insets (i), (ii), (iii) and (iv) are the same as the schematic regions shown in panel **b**. **d**, Side view of the calculated bending of the 18-mm long guidewire in the device region under the same applied force (F = 10 nN). Deflection of the tip of the 25-μm wide guidewire probe is 3 times that of the 75-μm wide guidewire probe (see Supplementary Text for calculation of the bending stiffness).

### Micro-endovascular probes and their bending properties

We first consider the design of MEV probes and how they meet the key constraints for implantation into the narrow and tortuous micron-scale brain vascular network without damaging the tissue. Inspired by minimally invasive catheter-based injection procedures (*19*), we consider designing polymer-based ultra-flexible MEV probes that can be loaded into and injected from flexible microcatheters. After inserting the microcatheter to the target vessel, the MEV probes can be injected into deeper vasculature by saline flow in the microcatheter. The microcatheter will then be retracted, leaving the probes in place. This procedure imposes several requirements on the probe design. First, the probes should be able to be loaded into and move smoothly in the microcatheter, controlled by saline flow. Second, the longitudinal bending stiffness of the MEV probes must be sufficient to ensure smooth injection into deeper vasculature beyond the reach of microcatheters without buckling, but also low enough to follow the tortuous vessels and prevent mechanical damage to vessel walls. Third, to optimize the neural recording signal quality, electrodes on the MEV probes should closely attach to the inner vessel walls (*15*).

The MEV probe (Fig. 1b and 1c) shows the ultra-flexible mesh-like device region on the left, tapering into the flexible stem in the middle, and then to the input/output (I/O) region on the right. Depending on the target region and species, the length of the stem can be varied. In this study, all MEV probes used are 7 cm in length. In the device region, metal electrodes are embedded in SU-8 polymer-based mesh-like substrate, which are delivered to the targeted brain region, while the I/O region remains exteriorized for subsequent connection and measurement. The injection depth can be determined by tracking the location of the I/O region during implantation.

Several aspects of the MEV probe structure have been considered. First, the transverse ribbons in the device region enable the 900-μm-wide probe to be rolled up inside the microcatheter, a flexible tube with an inner diameter of 200 μm and an outer diameter of 350 μm (*20*) (Fig. S1a). Second, a 10 μm thick guidewire made of SU-8 polymer (Fig. 1b, highlighted in red) was designed to optimize longitudinal bending stiffness for smooth injection into the tortuous vessel branches without buckling. The guidewire was placed over the middle ribbon (other longitudinal ribbons are 0.8 μm thick) in the device region, the stem and the I/O regions. The guidewire determines the overall bending stiffness of the MEV probe. The probes were manufactured, released from substrates, and loaded into microcatheters (*20-22*). With manual injection, the probes advance smoothly within the flexible microcatheter. The probes will remain extended without buckling even when the microcatheter is bent (Movie S1). Altering the guidewire width can modify the deflection of the probe tip under a given applied force (Fig. 1d, Supplementary Text). Third, the mesh-like structure relaxes and unfolds after injection (*20*) (Fig. S1b), allowing the electrodes to adhere against the inner vessel walls. The rest of the probe is subsequently injected from the microcatheter (Fig. S1c). Sixteen 80-μm platinum electrodes are distributed over a 1-cm length in the device region (Fig. S1b), allowing the probing from different brain regions.

### Branch-selective endovascular implantation

To demonstrate endovascular implantation, we exploited the established surgical procedure used for rodent stroke models, middle cerebral artery occlusion (MCAO) (*23*), without introducing an occlusion. The common carotid artery (CCA) bifurcates into the external carotid artery (ECA) and ICA in the neck. The ICA segment in the brain branches to form two major cerebral arteries, the MCA and the anterior cerebral artery (ACA), which overlay the cortex and the olfactory bulb, respectively. To perform MCAO, a filament is inserted into the ECA and threaded through the ICA until it occludes the MCA/ACA bifurcation. Similarly, a microcatheter loaded with an MEV probe and attached to a syringe can be inserted into the MCA/ACA bifurcation without occluding it and the probe subsequently injected (Fig. 2a, Fig. S2). While the microcatheter can only reach the MCA/ACA bifurcation, the saline flow in the microcatheter allows the probe to be carried much deeper into either MCA or ACA branches. Following injection, the microcatheter is retracted, leaving the MEV probe implanted inside the MCA or ACA.

**Figure 2.**
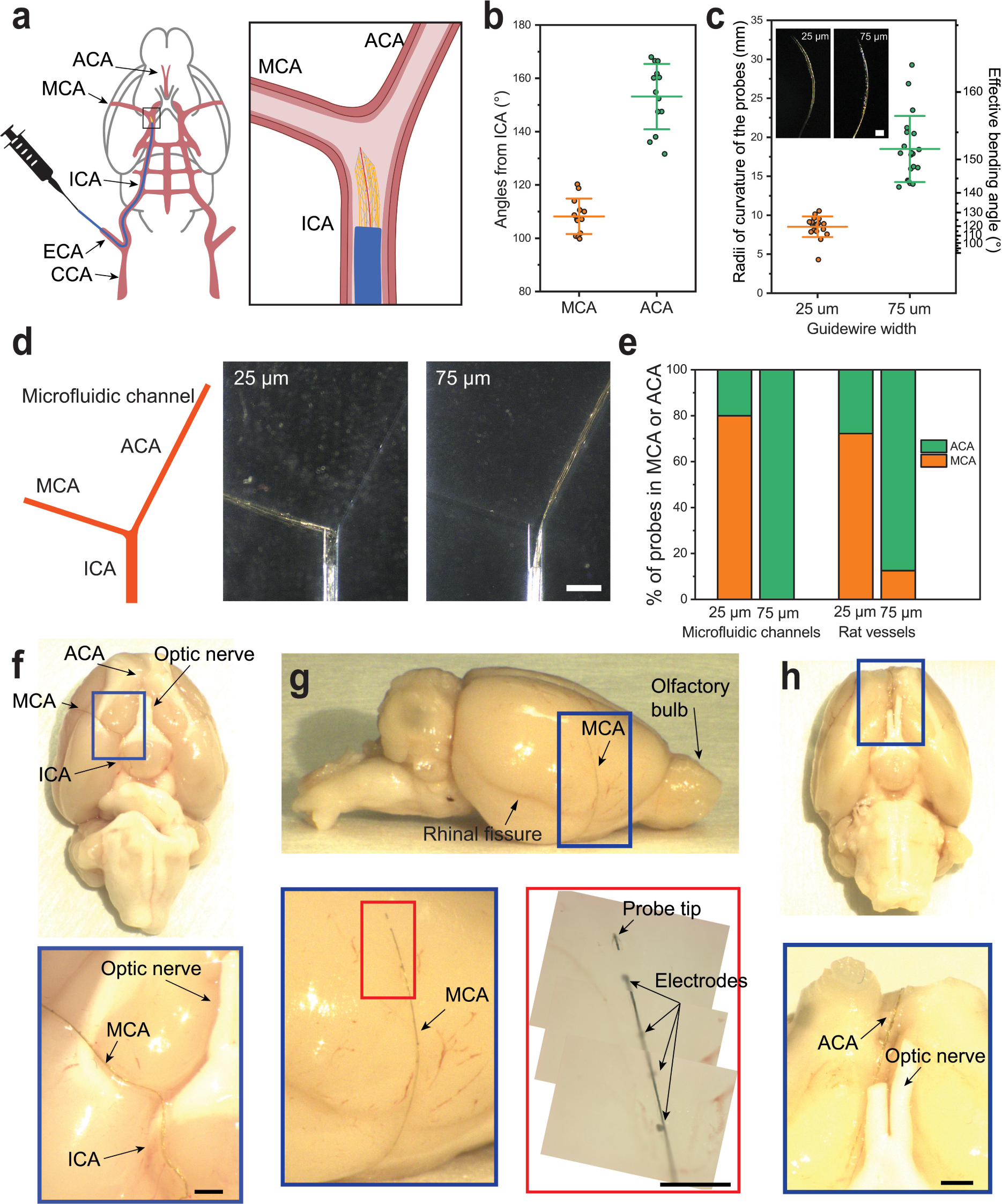
Branch-selective endovascular implantation. **a**, Schematics of endovascular implantation surgery. A microcatheter loaded with the MEV probe is inserted from the ECA opening to the MCA/ACA bifurcation. Saline injection carries the probe deeper into the MCA or ACA branch. **b**, Angles (N = 13) of the MCAs and the ACAs from the ICAs (with ±1 standard deviation, s.d.). **c**, Radii of curvature of the 18-mm long device region (N = 20, with ±1 standard deviation, s.d.) in water after release of the probes from the fabrication substrate. The effective bending angles are calculated based on the bending of the 18-mm long device region. Insets are representative images of the side views of the device regions of free-standing probes with 25-μm and 75-μm guidewires in water. Bending of the probes is due to the residue stress of SU-8 generated during fabrication. **d**, Left, schematic of a microfluidic channel designed to mimic the MCA/ACA bifurcation for testing branch-selective implantation. Right, representative implantation of a 25-μm guidewire probe into the MCA channel and a 75-μm guidewire probe into the ACA channel. Scale bar is 1 mm. **e**, Percentage of probes with different guidewire widths injected into MCA and ACA microfluidic channels (N = 5 for 25-μm guidewire probes, N = 11 for 75-μm guidewire probes) and rat vessels (N = 18 for 25-μm guidewire probes, N = 8 for 75- μm guidewire probes). **f-g**, Inferior view (**f**) and sagittal view (**g**) of a dissected and perfused rat brain with an MEV probe in the MCA. Zoom-in views show the probe maintains its elongated shape in the micron-scale vessels. The mark of the probe tip and Pt electrodes are visible through the vessel walls. **h**, Inferior view of a rat brain with a probe in the ACA. Scale bars in **f**-**h** are 1 mm.

Next, we achieve branch-selective implantation by tuning mechanical properties of the probe. Since the MCA and the ACA branches are at different angles from the ICA, we sought to determine whether changing the bending angle and bending stiffness of the guidewire would enable selective targeting of MCA and ACA branches. The angles of MCA and ACA from ICA (N = 13, each) were measured to be 108 ± 7° and 153 ± 12°, respectively (Fig. 2b). To match the bending angles of MCA and ACA, we fabricated MEV probes with guidewire widths of 25 μm and 75 μm. Due to the residual stress in the SU-8 film generated during processing (*24, 25*), the device region of the free-standing 25-μm and 75-μm guidewire probes formed radii of curvature of 8.5 ± 1.3 mm and 18.5 ± 4.2 mm in saline, respectively (N = 20) (Fig. 2c, Fig. S3), which correspond to effective bending angles of 119° and 152°. As the probe is injected through the microcatheter, it will encounter either the desired channel opening with the matching angle or the vessel wall. The operator can easily feel the difference: if the probe enters a vessel, the probe can continue to be injected, if it collides with the wall, they can retract the probe, move or rotate the catheter, and attempt again. To test the branch selectivity in vitro, we fabricated polydimethylsiloxane (PDMS) microfluidic channels matching the diameters and angles of the proximal segments of the ICA, the MCA and the ACA (Fig. 2d). Microcatheters carrying 25-μm and 75-μm guidewire MEV probes were inserted into the ICA channel, the probes were manually injected into either the MCA or the ACA channels.

An analysis of the branch selectivity (Fig. 2e) in microfluidic channels shows that 80% (N = 5) of the probes with 25-μm guidewires were injected into the MCA channel, while 100% (N = 11) of the probes with 75-μm guidewires were injected into the ACA channel. Similar branch selectivity was observed for the MEV probes implanted in rat brains *in vivo*: 72% (N = 18) of the probes with 25-μm guidewires were implanted into the MCAs, while 88% (N = 8) of the probes with 75-μm guidewires were implanted into the ACAs. The inferior/horizontal and sagittal views of the perfused and dissected rat brains confirmed the implantation into the MCA (Fig. 2f and 2g) and the ACA (Fig. 2h) branches, where the typical injection depth exceeds 1 cm into the distal segments pass the microcatheter opening at the MCA/ACA bifurcation. Magnified views of the brains with implanted probes (Fig. 2f-2h) reveal several important features. First, the MEV probes maintain their extended shapes within the distal segments of the targeted vessels without buckling. Second, the individual electrodes and the mark of the probe tip are easily identifiable. Third, the ultra-flexible device region is rolled up within the vessel, squeezing the 80-μm electrodes to the midline, indicating that the inner diameters of the targeted vessels are on the 100-micron scale.

### *In vivo* endovascular recording

With probes implanted into the MCA (Fig. 3a) and the ACA of anesthetized rats (Fig. S4), we demonstrated the ability of the MEV probes to record brain activity. Representative multichannel recordings (Fig. 3b, Fig. S5) yielded well-defined signals across all 16 channels. The fluctuation amplitude (200 µV-2 mV) and the dominant frequency (< 2 Hz) recorded are characteristic of the delta wave local field potentials under ketamine/xylazine anesthesia (*26*).

**Figure 3.**
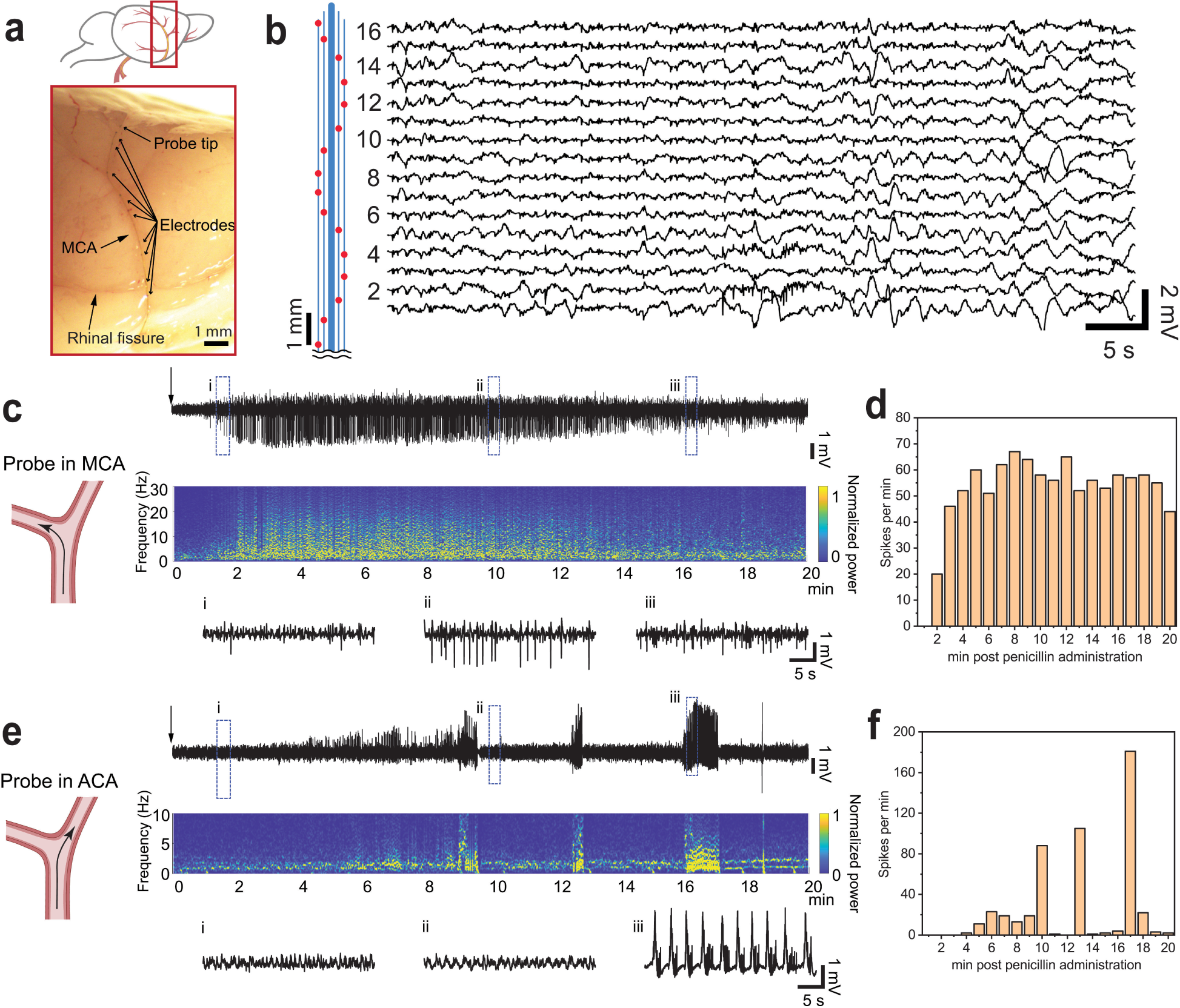
*In vivo* endovascular recording. **a-b**, Sagittal view of a rat brain showing an MEV probe in the MCA (**a**) and the corresponding acute *in vivo* 16-channel unfiltered recording showing local field potential oscillations under ketamine/xylazine anesthesia (**b**). The relative positions of 16 Pt electrodes are marked by red spots in the schematic (left panel in **b**), higher-numbered channels were implanted deeper in the MCA. **c**, Top, penicillin-induced seizures recorded by a representative channel from a probe in MCA for 20 min. Middle, spectrogram post penicillin injection. The colormap shows the normalized power levels. Bottom, zoom-in views showing the evolution of seizure spikes at three different time points. **ci**-**ciii** show the changes in spike amplitude over time. Black arrows denote the time point of penicillin injection. **d**, Number of spikes with amplitude over 1 mV per min recorded by the MCA probe in (**c**). 1034 spikes were recorded in 20 min. **e-f**, Recording and analysis of penicillin-induced seizures from a probe in ACA. **ei**-**eiii** show the recording trace before, between and during the burst firing activity. 496 spikes were recorded in 20 min in (**f**). The seizure spikes spread to the MCA region faster than ACA region. After seizure onset, the spikes recorded by the MCA probe reached constant frequency, while those recorded by the ACA probe showed fast spiking in waves every several minutes. The spikes in ACA region are consist of periodic oscillations followed by fast and narrow spikes.

Following this, we investigated whether the MEV probes implanted in the MCAs covering the cortex and the ACAs covering the olfactory bulb reveal different firing properties of different brain regions in neurological disease models. In anesthetized rats, we created epilepsy models by inducing local seizures with intracortical penicillin injection (*27, 28*) into the right hemisphere where the probes were implanted (Fig. 3c and 3e). Electrophysiological recordings by a representative channel on a probe implanted in the MCA (Fig. 3c) shows seizure activity characterized by bilateral spikes and spike-wave complexes. Following penicillin administration, seizures began almost immediately and reached a constant level after about 4 min (Fig. 3c and 3d). The mean spike frequency and amplitude from 5–20 min were 57 ± 6 / min and 1.95 ± 0.62 mV, respectively. Notably, simultaneous recording from all 16 channels of this probe identified spikes only in three adjacent channels (Fig. S6), demonstrating the ability of the MEV probes to locate and track the seizure foci. Recordings from MEV probes in MCAs in two other animals recorded similar firing patterns (Fig. S7). In comparison, representative recordings from a probe implanted in the ACA showed a latent phase lasting about 4 min after penicillin administration (Fig. 3e and 3f). After seizure onset, a burst-suppression pattern appeared with high-frequency spikes at around 10, 13, and 17 min. The burst firing activity around 17 min consists periodic field potential waves (frequency: 0.88 Hz, peak width: 95.8 ± 20.5 ms, amplitude: 2.89 ± 0.24 mV) followed by a train of fast and narrow spikes (frequency: 14.65 Hz, peak width: 1.9 ± 1.0 ms, amplitude: 0.71 ± 0.24 mV) (Fig. 3e). Seizure recordings from ACAs in two other animals showed similar firing patterns (Fig. S8).

Comparing the data recorded in the MCAs and the ACAs reveals several important characteristics. First, the seizure spikes spread to the cortex (MCA territory) faster than the olfactory bulb (ACA territory), suggesting that penicillin-induced seizures start as focal activity, then propagate to other brain regions, inducing generalized seizures (*29*). Second, the spike frequency, amplitude, shape and latent time in the cortex regions are also consistent with previous reports of rats treated with intracortical penicillin administration (*27, 28*). Third, the burst-suppression pattern recorded from the ACA probes was similar to that observed from rat olfactory bulb mitral cells (*30*). Fourth, the periodic field potential waves followed by spike trains also demonstrated the same firing behavior as the spontaneous activity in the olfactory bulb, in which slow oscillations are synchronized with the respiratory rhythm, followed by gamma bursts (fast spike trains) every respiratory cycle (*31*).

### Single unit activity recording

We then examine the possibility of recording single-unit spikes across the blood vessel wall, which has never been achieved using previous endovascular probes, due to their inability to target micron-scale vasculature with thin vessel walls. Typical depth electrodes detect spikes from neurons approximately 130 μm away (*32*). A 100-μm artery has a vessel wall thickness of approximately 10 - 20 μm (*33*), which is well within the detection range. Indeed, from the MEV probes injected deeply into the ACA segment overlaying the olfactory bulb under isoflurane anesthesia (Fig. 4a), we repeatedly recorded discontinuous, prolonged spontaneous bursts of action potentials with spindle shape lasting for tens of seconds (Fig. 4b), characteristic of olfactory bulb mitral cells of anesthetized rats (*30, 34*). The recording trace of the channels with the largest amplitude activity exhibits single-unit spikes. Single neuron activity was tracked by clustering the sorted spikes with principal component analysis from the channels with single unit activity (*21*), with three neurons identified from Ch. #1, and two neurons identified from Ch. #2 to Ch. #5 (Fig. 4c, top). All neurons exhibit higher spike amplitude and number of spikes in Ch. #1 and decaying from Ch. #2 through Ch. #5, indicating that Ch. #1 is the closest to the spiking neurons (Fig. 4c, bottom). Similar recordings were reproducibly observed from other MEV probes in ACAs (Fig. S9). In addition, from another MEV probe injected into the ACA under isoflurane anesthesia, we have observed burst spikes nested onto the respiratory rhythm in the field potential (*35, 36*) (Fig. 4d). The sharp downward spikes (Fig. 4e) showed a uniform potential waveform with average duration of ∼1 ms and peak-to-peak amplitude of ∼60 µV, characteristic of single-unit action potentials (*21*). We tested modulating neuron firing by raising isoflurane concentration (Fig. 4f) from 1.5% to 2%, which suppressed spiking. Decreasing to 0.5% restored it, which eventually disappeared, likely due to the prolonged exposure to the high concentration isoflurane (*37*).

**Figure 4.**
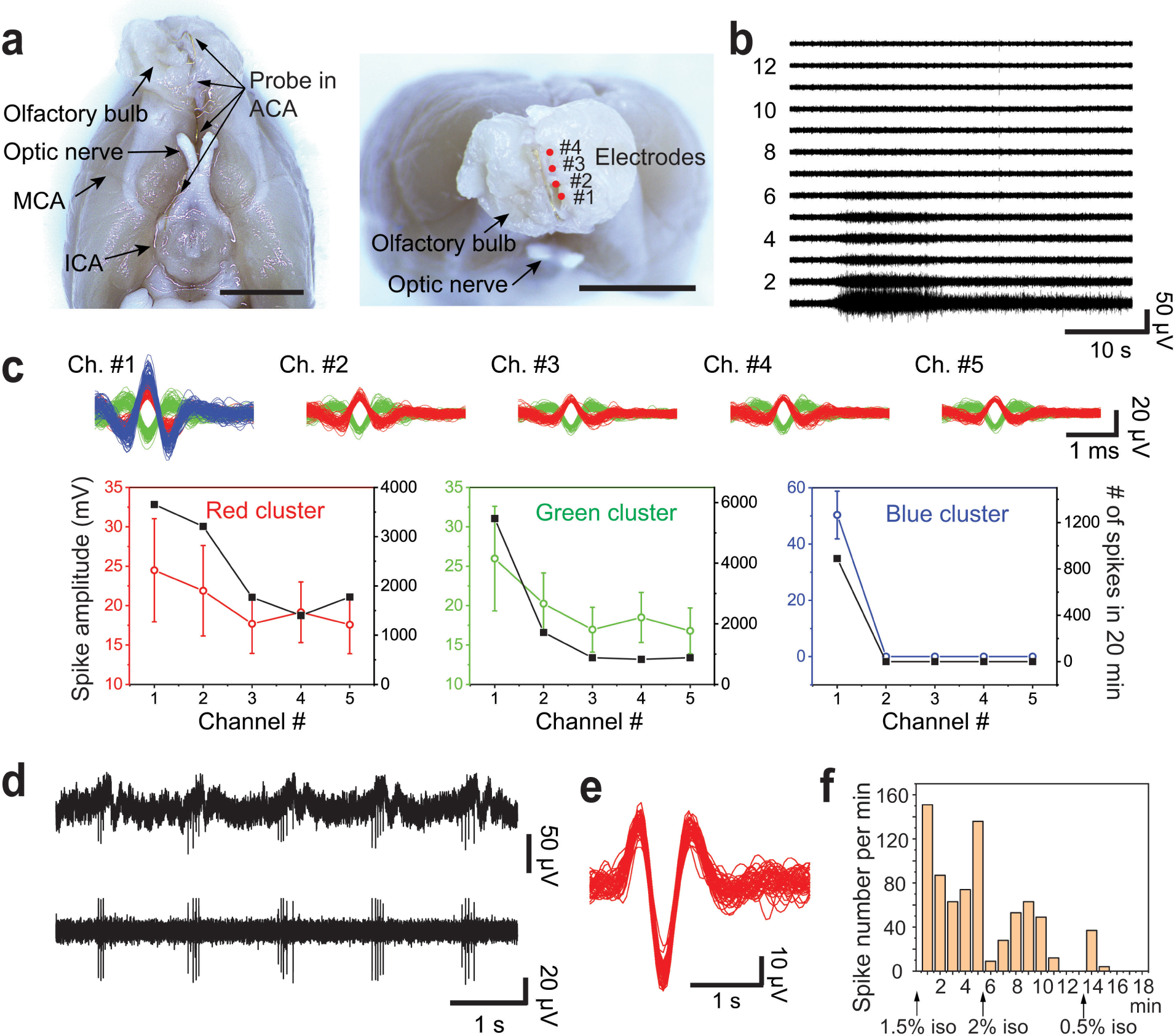
Single unit recording. **a-b**, Inferior (left) and anterior (right) views of a rat brain with an MEV probe in ACA in the olfactory bulb (**a**) and the corresponding multi-channel recording from the same probe after 250–6,000 Hz band-pass filtering to show single unit burst activity (**b**). Higher-numbered channels were implanted deeper in the ACA. Scale bars in **a** are 5 mm. **c**, Top, sorted spikes assigned to different neurons from the channels with single unit activity. Each distinct color in the sorted spikes represents a unique identified neuron. Bottom, spike amplitude (with ±1 standard deviation, s.d.) and the number of single unit spikes recorded in 20 min. **d**, Periodic single-unit spikes recorded by a probe in ACA. Top, the unfiltered recording trace. Bottom, after 250–6,000 Hz band-pass filtering. **e**, Single-unit spikes sorted from the data shown in **d**. **f**, Changes of firing frequency with different isoflurane concentration, where higher concentration (2.0%) decreased and eventually suppressed firing, while lower concentration (0.5%) temporarily recovered firing.

### Chronic histology

Next, we examined both the short-term and the chronic effects of endovascular implantation of MEV proves. For the short-term effects, laser doppler flowmetry was used to monitor the cerebral blood flow before and immediately after probe injection. Representative laser doppler flowmetry traces of probe implantation in the MCA and the ACA (Fig. S10) show that probe implantation does not have a significant effect on the cerebral blood flow. Immediately after MCA implantation, the blood flow fluctuated between 60% to 140% of the baseline, but the average value remains at 100%. The blood flow throughout the implantation processes remained much higher than the laser doppler flowmetry level required to induce stroke, which should be stabilized below 30% of the baseline for 90 min (*38*). On days 1, 3, 7, 14 and 28 after MEV probe implantation, we conducted behavior tests using neurological severity scores (NSS) (*39*). By day 3, all rats had reached a score of 0, indicating no neurologic deficit.

We then examine how chronic MEV probe implantation affects blood vessel walls and brain tissue. A histology study of the MCA was performed 28 days after implantation since neointimal formation reaches its maximum after rat artery stenting, thickening the walls substantially (*40, 41*), and the rat MCAO models start to recover from ischemic damage after 28 days (*42*). First, the blood–brain barrier (BBB) integrity was evaluated with immunoglobulin G (IgG) staining. The amount of IgG protein leaked into brain tissue after stroke is usually used to measure BBB leakage in rat brains (*43*). IgG-stained brain slices 28 days after implantation in MCA (Fig. 5a) showed no increase in IgG protein in the ipsilateral hemisphere when compared with the contralateral hemisphere, which confirms that the integrity of the BBB was well preserved. Second, cross-sections of the contralateral and ipsilateral MCAs were examined after Hematoxylin and Eosin (H&E) staining (Fig. 5b, Fig. S11). It was found that the probe ribbons were embedded in the vessel walls near the brain tissue. The vessel wall thicknesses of the contralateral and ipsilateral hemispheres are 18.9 ± 3.2 μm and 17.9 ± 2.0 μm measured from 15 brain slices from three rats (Fig. 5c, N = 5 from each rat, brain slices from the same rat are 600 μm apart). On the ipsilateral brain slices, no increase in vessel wall thickness was observed, confirming that MEV implants did not cause neointimal formation, which is commonly observed following vascular stenting (*40, 41*). Third, the lateral cerebral cortex within the MCA territory was evaluated with confocal fluorescence microscopy (Fig. 5d). Microglia, astrocytes, and nuclei were identified using antibodies against Iba1 (green), glial fibrillary acidic protein (GFAP, red), and DAPI (blue). The numbers of microglia counted from both hemispheres from 15 brain slices from three rats are 24 ± 6 on the contralateral side, and 24 ± 5 on the ipsilateral side, while the numbers of astrocytes are 53 ± 17 on the contralateral side, and 56 ± 14 on the ipsilateral side. There was no increase in microglia and astrocytes, indicating that endovascular implantation does not induce a significant immune response. Moreover, short-term histology studies were conducted 3 days post-implantation in the MCA (Fig. S12) and the ACA (Fig. S13). In both cases, there was no increase in vessel wall thickness or number of immune cells. Our chronic histology results showed a significant improvement from previously reported stiff endovascular probes, which could induce chronic venous thrombosis and occlusion (*15*). These observations not only demonstrated the minimal invasiveness of the MEV probes, but also indicate significant advantages in chronic electrophysiology recording, as the accumulation of glial scar tissue near the brain electrodes is known to cause electrode failure in clinically relevant chronic settings (*44*).

**Figure 5.**
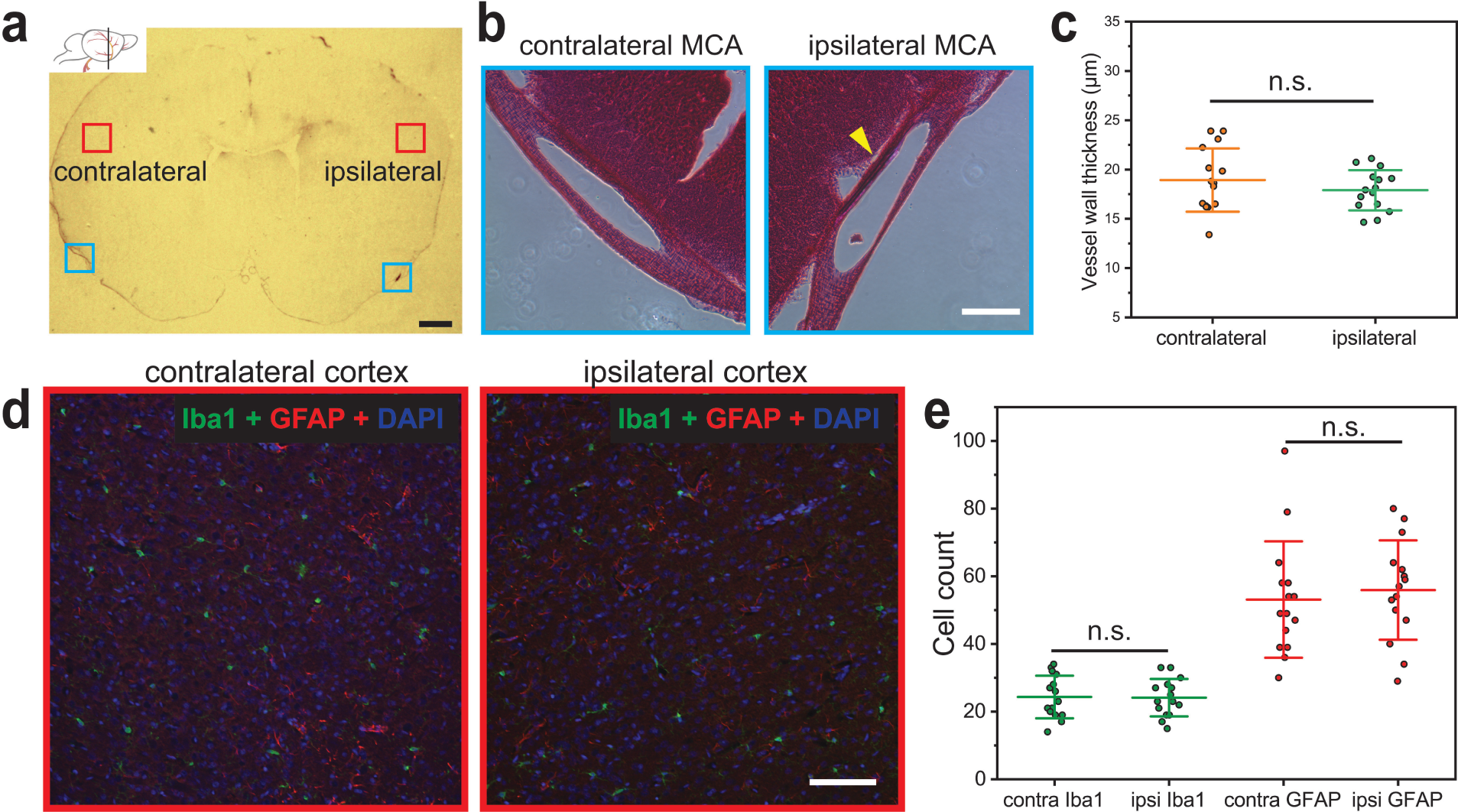
Chronic histology. **a**, Digital camera image of a representative IgG-stained coronal brain slice 28 days post-implantation in MCA. Scale bar is 1 mm. Inset shows the location of the slice. **b**, Representative zoom-in views of the contralateral and ipsilateral MCA cross-sections from the H&E-stained slice from the regions highlighted by the blue boxes in **a**. The yellow arrow highlights the probe embedded in the vessel wall. Scale bar is 100 μm. Note that the vessels were cut at an angle not perpendicular to the central axis, accordingly, they have an oval cross-section, the vessel diameter is the minor axis. **c**. MCA vessel wall thickness measured from H&E images of 15 brain slices from three rats (N = 5 for each rat, brain slices from the same rat are 600 μm apart, with ±1 standard deviation, s.d.). **d**, Representative confocal fluorescence microscopy images of the contralateral and ipsilateral cortexes from the regions highlighted by the red boxes in **a**. The brain slice was stained with antibodies for Iba1 (green) and GFAP (red), and DAPI (blue). Scale bar is 100 μm. **e**, Number of microglia (Iba1) and astrocytes (GFAP) counted from fluorescence images of 600 μm*450 μm from 15 brain slices from three rats (N = 5 for each rat, brain slices from the same rat are 600 μm apart, with ±1 standard deviation, s.d.). n.s.= nonsignificant; unpaired two-tailed t test.

### Outlook

This study developed a method for delivering ultra-flexible electronics into 100-micron scale vessels in rodents without the need for open skull surgery. The MEV probe can be selectively implanted into small vessel branches that are not accessible to any available microcatheters, thus enabling neural recording across vessel walls at single cell resolution. Histology studies of the probe/vessel wall/brain tissue interface showed minimal immune response and long-term stability.

We anticipate that our approach will be applicable in a variety of other areas in the future. First, as MEV implantation induces minimal immune response, future efforts will be directed toward developing stretchable MEV probes with chronic recording capabilities. Second, the platform could be scaled to hundreds to thousands of electrodes for high-density recording and stimulation (*16, 45*) by adapting highly scalable I/O interfaces (*46*). Third, steering compartments, such as magnetic actuators (*18, 47*), could be incorporated to the tip of the probe to control navigation and improve the branch selectivity. Finally, the element size on the probes could be further reduced (*22*) to target the capillary network. This platform technology can be extended to the detection and treatment of many neurological diseases as a research tool, and serve as the foundation for future clinical translation of minimally invasive neuroelectronic interfaces.

## Acknowledgements

The authors gratefully acknowledge Professor Zhenan Bao at Stanford University and Dr. Theodore J. Zwang at Massachusetts General Hospital for helpful discussions. C.M.L. acknowledges support from Air Force Office of Scientific Research (FA9550-18-1-0469, FA9550- 19-1-0246) and National Institutes of Health Director’s Pioneer Award (DP1EB025835). E.H.L. acknowledges support from National Institutes of Health (R01NS099620). C.M.S. acknowledges support from National Institutes of Health (R01NS107445). A.Z. acknowledges support from American Heart Association (23POST1018301). This work was performed in part at the Harvard University Center for Nanoscale Systems (CNS); a member of the National Nanotechnology Coordinated Infrastructure Network (NNCI), which is supported by the National Science Foundation under NSF award no. ECCS-2025158. This work was performed in part in the nano@Stanford labs, which are supported by the National Science Foundation as part of the National Nanotechnology Coordinated Infrastructure under award ECCS-2026822.

## Author contributions

A.Z. and C.M.L. conceived and designed the experiments, A.Z., E.T.M. and L.X. performed the experiments, A.Z. and C.M.L. and analyzed the data and wrote the paper. All authors discussed the results and commented on the manuscript.

## Additional information

Supplementary information is available in the online version of the paper. Reprints and permissions information is available online at www.nature.com/reprints. Correspondence and requests for materials should be addressed to A.Z., E.H.L., and C.M.L.

## Data availability

The data that support the findings of this study are presented in the main text and the Supplementary Information, and are available from the corresponding authors on reasonable request.

## Methods

### Design and fabrication of MEV probes

MEV probes with double-sided electrodes and input/output (I/O) metal regions (*48*) are fabricated using standard photolithography. (1) A 100-nm-thick Ni sacrificial layer was thermally evaporated (Sharon Vacuum Co.) onto a 4-inch Si wafer (n-type 0.005 Ωcm, 1,000 nm thermal oxide, NOVA Electronic Materials). (2) LOR 3A and S1805 (Microchem) were spin-coated at 4,000 rpm for 45 s and baked at 180 °C for 4 min and at 115 °C for 1 min, respectively. The photoresist was patterned by photolithography with a mask aligner (SUSS MA6 mask aligner, SUSS MicroTec) and developed (MF-CD-26, MicroChem Corp.) for 1 min. Following this photolithography process, a 100-nm-thick Au bottom metal input/output (I/O) layer was deposited by thermal evaporation (Sharon Vacuum Co.), followed by a liftoff step (Remover PG, MicroChem). (3) The photolithography process in step 2 was repeated to define 80-μm Pt bottom electrode regions for E-beam evaporation (Denton Vacuum Co.), followed by a liftoff step. (4) For the bottom-passivation layer, negative photoresist SU-8 (SU-8 2000.5; MicroChem) was spin-coated on the Si wafer at 3,000 rpm for 30 s, pre-baked at 65 °C for 1 min and 95 °C for 1 min, and then patterned by photolithography with the mask aligner. After post-baking at 65 °C for 1 min and 95 °C for 1 min, The SU-8 resist was developed in SU-8 developer (Microchem) for 1.5 min, rinsed with isopropanol, dried in N_2_ and hard baked at 185 °C for 1 h. (5) The photolithography process in step 2 was repeated to define metal interconnects, followed by 100- nm-thick Au deposition by thermal evaporation and liftoff. (6) The top passivation layer is then patterned using a similar procedure as in step 4, followed by hard baking at 195 °C for 1 h. (7) The process in 3 was repeated to deposit the top layer of Pt electrodes. (8) For the ∼10-μm guide wire layer, negative photoresist SU-8 (SU-8 2010; MicroChem) was spin-coated on the Si wafer at 3,000 rpm for 30 s, pre-baked at 65 °C for 1 min and 95 °C for 2 min, and then patterned by photolithography with the mask aligner. After post-baking at 65 °C for 1 min and 95 °C for 2.5 min, The SU-8 resist was developed in SU-8 developer (Microchem) for 3 min, rinsed with isopropanol, dried in N_2_ and hard baked at 200 °C for 2 h. (9) To release the probes, the Si wafer was cleaned with oxygen plasma (75 W, 1 min) and immersed in a Ni etchant solution comprising 40% FeCl_3_:39% HCl:H_2_O=1:1:10 on a 50 °C hot plate for 60 min. Released probes were rinsed with deionized (DI) water for three times and sterilized as described in the next section. (10) Before implantation, the probe was loaded into a flexible microcatheter. The microcatheter was prepared by fixing a 30-cm long PE-8 tube (ID = 200 μm, OD = 355 μm, SAI Infusion) on a 23-G needle and sealed with UV light glue (Visbella) and cured with handheld UV flashlight (Vansky). A 5- ml syringe filled with sterile saline was attached to the 16-G needle and infused the PE-8 microcatheter with saline. The end of the tube was positioned near the I/O pads of the probe, the syringe was manually retracted to draw the probe into the microcatheter. The 5-ml syringe was replaced with a 1-ml syringe filled with saline before implantation.

### Fabrication of PDMS microfluidic channels

To fabricate the SU8 mold, SU8 2150 (Microchem) were spin-coated at 500 rpm for 5s at a ramp rate of 100 rpm/s and 1,400 rpm for 30s at a ramp rate of 300 rpm/s, and baked at 65 °C for 10 min and 95 °C for 2 h. The photoresist was patterned by photolithography with a mask aligner (SUSS MA6 mask aligner, SUSS MicroTec), developed in SU-8 developer (Microchem) for 45 min, and hard baked at 180 °C for 1 h. The thickness of SU-8 layer after hard bake is about 350 μm. The width of the ICA channel is 400 μm and the MCA and the ACA channels are 200 μm. After hard bake, the substrates were coated with hexamethyldisilazane (HDMS) in YES oven. To create PDMS channels, liquid PDMS prepolymer was mixed at a 10:1 ratio with the curing agent and poured onto the SU8 mold. The PDMS was cured at 70 °C for 2 h, peeled off the mold, treated with oxygen plasma at 50 W for 1 min (March Instruments PX-250), and attached to a 4- inch Si wafer.

### *In vivo* endovascular implantation surgery

Adult (300-400 g) male Wistar rats (Charles River Laboratory) were used in the study. All procedures were approved by the Institutional Animal Care and Use Committee (IACUC) at Massachusetts General Hospital (MGH) and Administrative Panel on Laboratory Animal Care (APLAC) at Stanford University. The animal care and use programs at MGH and Stanford meet the requirements of federal law (89–544 and 91–579) and NIH regulations and are also accredited by the American Association for Accreditation of Laboratory Animal Care (AAALAC). Before surgical procedures, animals were group-housed on a 12h:12h light:dark schedule and fed with food and water ad libitum. Animals were housed individually after surgical procedures.

*In vivo* implantation of the MEV probes into vessels of rat brains was performed using manually controlled injection. All metal tools in direct contact with the animal subjects were bead-sterilized (Fine Science Tools) for 1 h before use, and all plastic tools in direct contact with the animal subjects were sterilized with 70% ethanol and rinsed with sterile DI water and sterile saline before use. The MEV probes were sterilized with 70% ethanol followed by rinsing in sterile DI water and sterile saline before injection. Rats were anaesthetized either under 100 mg kg^−1^ of ketamine and 20 mg kg^−1^ xylazine (Patterson Veterinary Supply), or spontaneous respiration with isoflurane (3% for induction, 1.5% for maintenance, Matrx vaporizer) in 30/70% oxygen/nitrous-oxide mixture. The degree of anesthesia was verified via toe pinch before surgery. Rectal temperature was maintained at 37.5 °C with a thermostat-controlled heating pad (EZ-TC-1000-M Homeothermic Temperature Controller). Puralube vet ointment (Dechra Pharmaceuticals) was applied on both eyes to prevent corneal damage. Hair removal lotion (Nair, Church & Dwight) was applied to the scalp for depilation and an alternating series of Betadine surgical scrub (Purdue Products) and 70% alcohol was applied to sterilize the depilated scalp skin. A sterile scalpel was used to make a 5 mm longitudinal incision in the scalp along the sagittal sinus. The scalp skin was resected to expose a 5 mm×5 mm portion of the skull. A 1-mm-diameter burr hole was made using a dental drill (Micromotor with On/Off Pedal 110/220, Grobet USA) according to the following stereotaxic coordinates: anteroposterior, −3 mm; mediolateral, −3 mm. A 3-cm long sterilized stainless steel wire (0.015”, Malin Company) was bent and ∼0.5 mm at one end was inserted into this burr hole and fixed with Metabond adhesive cement (Parkell) to serve as the grounding and reference electrode. A flat, flexible cable (FFC, WM11484-ND, Digi-Key Electronics) with 32 conductors with a 0.50-mm pitch was attached onto the skull with Metabond adhesive cement (Fig. S4). For seizure recording and laser doppler flowmetry recording that required access to the skull, the FFC was placed next to the surgery region.

The ventral neck region was then depilated and sterilized. The rat was placed under a dissection microscope (LEICA MZ75), and a pair of sterile scissors (Fine Science Tools) was used to make a 2 cm midline incision in the ventral neck region. Retractors (Braintree Scientific) were used to separate the muscles and expose the right common carotid artery (CCA), external carotid artery (ECA), and internal carotid artery (ICA). The arteries were then detached from surrounding tissues. On ECA, the superior thyroid artery (STA) branch and occipital artery (OA) branch were coagulated with a bipolar coagulator (Kirwan Surgical Products), and cut at the coagulated segment.

4-0 silk sutures (Fine Science Tools) were used to ligate the ECA and temporarily stop blood flow in CCA and posterior auricular artery (PAA). A vascular clamp (Fine Science Tools) was applied to ICA. A small incision was then made at the distal end of the ECA stump with Vannas spring scissors (Fine Science Tools). Next, a subcutaneous tunnel was made from the neck incision to the FFC on the skull. The microcatheter loaded with an MEV probe was threaded through the subcutaneous tunnel and the tip was inserted into the ECA stump (Fig. S2a). A loose suture was placed on the ECA stump to prevent blood leakage around the microcatheter. The vascular clamp on ICA was removed, and the microcatheter tip was advanced into ICA by 15-18 mm until reaching the bifurcation of middle cerebral artery (MCA) and ICA. The probe was manually injected into the deeper branches by pushing the plunger. As the probe is injected through the microcatheter, it will encounter either the desired channel opening with the matching angle or the vessel wall. The operator can easily feel the difference: if the probe enters a vessel, the probe can continue to be injected, if it collides with the wall, they can retract the probe, move or rotate the catheter, and attempt again. The implantation depth is tracked by the location of the I/O pads in the microcatheter. The microcatheter was then retracted slowly (Fig. S2b) from the subcutaneous tunnel, leaving the probe stem under the skin. The ECA was then ligated with suture to fix the probe stem at the ECA opening. The vascular clamp and sutures on ICA, CCA and PAA were removed to allow reperfusion. The I/O pads on the probe were then aligned with flowing DI water onto the conductors of the FFC (Fig. S4), and dried with an air duster spray (VWR International). The incision in the neck region was closed by silk sutures.

After surgery, the rats for acute recording under anesthesia were transcardially perfused with PBS and 4% paraformaldehyde, while the rats for chronic implantation were returned to cages placed on a 37 °C heating pad. The activity of the rats was monitored until it was fully recovered from anesthesia. Buprenex (Buprenorphine, Patterson Veterinary Supply) analgesia was given intraperitoneally at a dose of 0.05 mg per kg body weight every 12 h for up to 72 h post-surgery.

### Electrophysiology recording

The electrophysiology of rats with implanted MEV probes was recorded with the FFC connected to an Intan RHD 2132 amplifier evaluation system (Intan Technologies). Electrophysiological recordings were acquired with a 20 kHz sampling rate and a 60 Hz notch filter.

To induce local seizures, before probe implantation, penicillin solution (1,000-2,000 units / 50 μL, Sigma-Aldrich) was loaded into a 1-ml syringe with a 30-G needle. The needle was bent and inserted into the cortex (anteroposterior, −2 mm; mediolateral, 2 mm; dorsoventral, 1.50 mm) and fixed on the skull with Metabond adhesive cement. During electrophysiological recording, the penicillin solution was injected by pushing the plunger. After recording for up to 30 min, rats were euthanized by cardiac perfusion.

### Laser Doppler Flowmetry measurement

Laser Doppler Flowmetry (PeriFlux System 5000) was used to evaluate cerebral blood flow before and right after probe injection. First, a 1.2 cm long midline incision in the scalp to expose the skull bone. After the tissues and muscle on right side of the skull were removed, a 1.5 mm diameter dimple was drilled according to the following stereotaxic coordinates: anteroposterior, 0 mm; mediolateral, 5 mm. The laser doppler flowmetry probe was fixed on the dimple with Krazy glue to monitor the blood flow. The flowmetry data was acquired with LabChart software (AD Instruments).

### Histology sample preparation, immunostaining, and imaging

Rats implanted with MEV probes were anaesthetized with 3% isoflurane or 300 mg kg^−1^ of ketamine (Patterson Veterinary Supply) and transcardially perfused with ice-cold 30 ml saline and 30 ml 4% paraformaldehyde (PFA, Electron Microscopy Sciences) at specified times post-injection, followed by decapitation. The scalp skin was cut and the exposed skull was removed by bone cutters (Fine Science Tools). The brain was dissected from the bottom of the skull by cutting the connecting tissues and blood vessels, and was stored in 4% PFA for 1-2 days at 4 °C. The brain was then transferred to 15% sucrose solution (Sigma-Aldrich) in saline for 1 day until the brain sinks and then stored in 30% sucrose solution in saline for 1 day at 4 °C until the brain sinks.

The brain was trimmed and placed in optimum cutting temperature (OCT) compound (Tissue-Tek) in a cryomold cup (Tissue-Tek) for 30 min. The cryomold cup was then placed in 2- methylbutane (Sigma-Aldrich) in dry ice until the OCT turns white. The frozen OCT block containing the brain was then glued on a metal holder in dry ice with more OCT compound. The metal holder was then placed in cryostat (Leica Biosystems) for at least 1 h. The brain was then cut into 20 μm and 50 μm slices and transferred onto glass slides (Sigma-Aldrich).

For Iba1 and GFAP staining, 20 μm brain slices were fixed in 4% PFA for 10 min, followed by rinsing in PBS for 3 times. The slices were then sequentially incubated with 0.5% Triton X-100 for 10 min, 3% BSA for 1 h, and 1:200 primary antibodies, anti-Iba1 (FUJIFILM Wako Pure Chemical Corp.) or anti-GFAP (Thermo Fisher Scientific) overnight, followed by rinsing in PBS for 3 times. The slices were then incubated with secondary antibodies (Vector Laboratories) labeled with Cy5 and Cy3, respectively, for 1 h and rinsed with PBS for 3 times. Finally, the slices were mounted with ProLong™ Gold Antifade Mountant with DAPI (Thermo Fisher Scientific).

For IgG staining, 20 μm brain slices were fixed in 4% PFA for 10 min, followed by rinsing in PBS for 3 times. The endogenous peroxidase activity was inhibited by incubating in peroxidase suppressor (Thermo Fisher Scientific) for 30 min and rinsed with PBS for 3 times. The slices were then sequentially incubated with 3% BSA for 1 h, and anti-Rat IgG (Thermo Fisher Scientific) overnight, followed by rinsing in PBS for 3 times. The slices were then stained with DAB Peroxidase (HRP) Substrate Kit (Vector Laboratories). Finally, the brain slices were dehydrated with 70% ethanol for 2 min twice, and 100% ethanol for 2 min twice, and mounted with coverslips. For Hematoxylin and Eosin (H&E) staining, 50 μm brain slices were labeled with Hematoxylin and Eosin Stain Kit (Vector Laboratories).

Fluorescence images were acquired on an Olympus FV1000 upright confocal microscope (BX61). 405, 559, and 635 nm laser sources were used with water immersion 20× objective as the excitation sources for DAPI, Cy3 and Cy5, respectively. All fluorescence images were acquired by Olympus Fluoview software v2.1.

Digital camera images of IgG-stained slices were acquired with an eyepiece camera (DCC1240C, Thorlabs Inc.) equipped with a manual zoom lens (MLH-10X, CBC Group), images were taken with ThorCam uc480 image acquisition software.

Optical images of IgG- and H&E-stained slices were acquired with a Nikon inverted microscope (Eclipse Ti-S) equipped with a QImaging camera and NIS-Elements Imaging Software.

### Behavior tests

Post-implantation, rats were evaluated using neurological severity scores (NSS) (*39*) on days 1, 3, 7, 14 and 28. NSS is a test involve motor, sensory, reflex and balance, on a scale of 0 to 18. 1 point is scored for the inability to perform each specific test, the higher score, the more severe is the injury.

## References

1. N. G. Hatsopoulos, J. P. Donoghue, The Science of Neural Interface Systems. Annu. Rev. Neurosci. 32, 249–266 (2009).

2. D. O. Adewole, M. D. Serruya, J. A. Wolf, D. K. Cullen, Bioactive neuroelectronic interfaces. Front. Neurosci. 13, 269 (2019).

3. J. Wolpaw, E. W. Wolpaw, Brain-computer interfaces: principles and practice. (Oxford University Press, 2012).

4. V. Sakkalis, Modern electroencephalographic assessment techniques. (Springer, 2015).

5. M. Hirata, T. Yoshimine, in Clinical Systems Neuroscience, Kansaku K., Cohen L. G., Birbaumer N., Eds. (Springer Japan, Tokyo, 2015), pp. 83-100.

6. G. Hong, C. M. Lieber, Novel electrode technologies for neural recordings. Nat. Rev. Neurosci. 20, 330–345 (2019).

7. L. Luan, J. T. Robinson, B. Aazhang, T. Chi, K. Yang, X. Li, H. Rathore, A. Singer, S. Yellapantula, Y. Fan, Z. Yu, C. Xie, Recent advances in electrical neural interface engineering: Minimal invasiveness, longevity, and scalability. Neuron 108, 302–321 (2020).

8. E. Hedegärd, J. Bjellvi, A. Edelvik, B. Rydenhag, R. Flink, K. Malmgren, Complications to invasive epilepsy surgery workup with subdural and depth electrodes: a prospective population-based observational study. J. Neurol. Neurosurg. Psychiatry 85, 716–720 (2014).

9. I. G. Gould, P. Tsai, D. Kleinfeld, A. Linninger, The capillary bed offers the largest hemodynamic resistance to the cortical blood supply. J. Cereb. Blood Flow Metab. 37, 52–68 (2017).

10. P. S. Tsai, J. P. Kaufhold, P. Blinder, B. Friedman, P. J. Drew, H. J. Karten, P. D. Lyden, D. Kleinfeld, Correlations of neuronal and microvascular densities in murine cortex revealed by direct counting and colocalization of nuclei and vessels. J. Neurosci. 29, 14553–14570 (2009).

11. R. K. Sefcik, N. L. Opie, S. E. John, C. P. Kellner, J. Mocco, T. J. Oxley, The evolution of endovascular electroencephalography: historical perspective and future applications. Neurosurg. Focus 40, E7 (2016).

12. J. Z. Fan, V. Lopez-Rivera, S. A. Sheth, Over the horizon: The present and future of endovascular neural recording and stimulation. Front. Neurosci. 14, (2020).

13. R. D. Penn, S. K. Hilal, W. J. Michelsen, E. S. Goldensohn, J. Driller, Intravascular intracranial EEG recording. J. Neurosurg. 38, 239–243 (1973).

14. J. M. Serrador, P. A. Picot, B. K. Rutt, J. K. Shoemaker, R. L. Bondar, MRI measures of middle cerebral artery diameter in conscious humans during simulated orthostasis. Stroke 31, 1672–1678 (2000).

15. T. J. Oxley, N. L. Opie, S. E. John, G. S. Rind, S. M. Ronayne, T. L. Wheeler, J. W. Judy, A. J. McDonald, A. Dornom, T. J. H. Lovell, C. Steward, D. J. Garrett, B. A. Moffat, E. H. Lui, N. Yassi, B. C. V. Campbell, Y. T. Wong, K. E. Fox, E. S. Nurse, I. E. Bennett, S. H. Bauquier, K. A. Liyanage, N. R. van der Nagel, P. Perucca, A. Ahnood, K. P. Gill, B. Yan, L. Churilov, C. R. French, P. M. Desmond, M. K. Horne, L. Kiers, S. Prawer, S. M. Davis, A. N. Burkitt, P. J. Mitchell, D. B. Grayden, C. N. May, T. J. O’Brien, Minimally invasive endovascular stent-electrode array for high-fidelity, chronic recordings of cortical neural activity. Nat. Biotechnol. 34, 320 (2016).

16. N. L. Opie, S. E. John, G. S. Rind, S. M. Ronayne, Y. T. Wong, G. Gerboni, P. E. Yoo, T. J. H. Lovell, T. C. M. Scordas, S. L. Wilson, A. Dornom, T. Vale, T. J. O’Brien, D. B. Grayden, C. N. May, T. J. Oxley, Focal stimulation of the sheep motor cortex with a chronically implanted minimally invasive electrode array mounted on an endovascular stent. *Nat*. Biomed. Eng. 2, 907–914 (2018).

17. H. Watanabe, H. Takahashi, M. Nakao, K. Walton, R. R. Llinás, Intravascular neural interface with nanowire electrode. Electron. Commun. Jpn. 92, 29–37 (2009).

18. L. Pancaldi, P. Dirix, A. Fanelli, A. M. Lima, N. Stergiopulos, P. J. Mosimann, D. Ghezzi, M. S. Sakar, Flow driven robotic navigation of microengineered endovascular probes. Nat. Commun. 11, 6356 (2020).

19. W. Sherman, T. P. Martens, J. F. Viles-Gonzalez, T. Siminiak, Catheter-based delivery of cells to the heart. Nat. Clin. Pract. Cardiovasc. Med. 3, S57–S64 (2006).

20. J. Liu, T.-M. Fu, Z. Cheng, G. Hong, T. Zhou, L. Jin, M. Duvvuri, Z. Jiang, P. Kruskal, C. Xie, Z. Suo, Y. Fang, C. M. Lieber, Syringe-injectable electronics. Nat. Nanotechnol. 10, 629–636 (2015).

21. T.-M. Fu, G. Hong, T. Zhou, T. G. Schuhmann, R. D. Viveros, C. M. Lieber, Stable long-term chronic brain mapping at the single-neuron level. Nat. Methods 13, 875–882 (2016).

22. X. Yang, T. Zhou, T. J. Zwang, G. Hong, Y. Zhao, R. D. Viveros, T.-M. Fu, T. Gao, C. M. Lieber, Bioinspired neuron-like electronics. Nat. Mater. 18, 510–517 (2019).

23. L. Belayev, O. F. Alonso, R. Busto, W. Zhao, M. D. Ginsberg, Middle cerebral artery occlusion in the rat by intraluminal suture. Neurological and pathological evaluation of an improved model. Stroke 27, 1616–1623 (1996).

24. S. Keller, D. Haefliger, A. Boisen, Fabrication of thin SU-8 cantilevers: initial bending, release and time stability. J. Micromech. Microeng 20, 045024 (2010).

25. S. Keller, G. Blagoi, M. Lillemose, D. Haefliger, A. Boisen, Processing of thin SU-8 films. J. Micromech. Microeng 18, 125020 (2008).

26. S. Chauvette, S. Crochet, M. Volgushev, I. Timofeev, Properties of slow oscillation during slow-wave sleep and anesthesia in cats. J. Neurosci. 31, 14998–15008 (2011).

27. M. Ayyildiz, S. Coskun, M. Yildirim, E. Agar, The effects of ascorbic acid on penicillin-induced epileptiform activity in rats. Epilepsia 48, 1388–1395 (2007).

28. M. Yildirim, C. Marangoz, Anticonvulsant effects of focal and intracerebroventricular adenosine on penicillin-induced epileptiform activity in rats. Brain Res. 1127, 193–200 (2007).

28. I. Akdogan, N. G. Yonguc, in Underlying mechanisms of epilepsy, F. S. Kaneez, Ed. (InTech, 2011), pp. 269–283.

30. G. Leng, H. Hashimoto, C. Tsuji, N. Sabatier, M. Ludwig, Discharge patterning in rat olfactory bulb mitral cells in vivo. Physiol. Rep. 2, e12021 (2014).

31. C. Martin, D. Houitte, M. Guillermier, F. Petit, G. Bonvento, H. Gurden, Alteration of sensory-evoked metabolic and oscillatory activities in the olfactory bulb of GLAST- deficient mice. Front. Neural Circuits 6, (2012).

32. A. Marblestone, B. Zamft, Y. Maguire, M. Shapiro, T. Cybulski, J. Glaser, D. Amodei, P. B. Stranges, R. Kalhor, D. Dalrymple, D. Seo, E. Alon, M. Maharbiz, J. Carmena, J. Rabaey, E. Boyden, G. Church, K. Kording, Physical principles for scalable neural recording. Front. Comput. Neurosc. 7, (2013).

33. M. J. Lew, J. A. Angus, Wall thickness to lumen diameter ratios of arteries from SHR and WKY: Comparison of pressurised and wire-mounted preparations. J. Vasc. Res. 29, 435–442 (1992).

34. G. Z. Yu, H. Kaba, H. Saito, K. Seto, Heterogeneous characteristics of mitral cells in the rat olfactory bulb. Brain Res. Bull. 31, 701–706 (1993).

35. B. N. Cazakoff, B. Y. B. Lau, K. L. Crump, H. S. Demmer, S. D. Shea, Broadly tuned and respiration-independent inhibition in the olfactory bulb of awake mice. Nat. Neurosci. 17, 569–576 (2014).

36. R. C. Gerkin, S. J. Tripathy, N. N. Urban, Origins of correlated spiking in the mammalian olfactory bulb. Proc. Natl. Acad. Sci. USA 110, 17083–17088 (2013).

37. N. Fujiwara, H. Higashi, S. Nishi, K. Shimoji, S. Sugita, M. Yoshimura, Changes in spontaneous firing patterns of rat hippocampal neurones induced by volatile anaesthetics. J. Physiol. 402, 155–175 (1988).

38. R. Schmid-Elsaesser, S. Zausinger, E. Hungerhuber, A. Baethmann, H.-J. Reulen, A critical reevaluation of the intraluminal thread model of focal cerebral ischemia. Stroke 29, 2162–70 (1998).

39. J. Chen, Y. Li, L. Wang, Z. Zhang, D. Lu, M. Lu, M. Chopp, Therapeutic benefit of intravenous administration of bone marrow stromal cells after cerebral ischemia in rats. Stroke 32, 1005–1011 (2001).

40. B. Langeveld, A. J. M. Roks, R. A. Tio, A. J. van Boven, J. J. L. van der Want, R. H. Henning, H. M. M. van Beusekom, W. J. van der Giessen, F. Zijlstra, W. H. van Gilst, Rat abdominal aorta stenting: A new and reliable small animal model for in-stent restenosis. J. Vasc. Res. 41, 377–386 (2004).

41. A. V. Finn, H. K. Gold, A. Tang, D. K. Weber, T. N. Wight, A. Clermont, R. Virmani, F. D. Kolodgie, A novel rat model of carotid artery stenting for the understanding of restenosis in metabolic diseases. J. Vasc. Res. 39, 414–425 (2002).

42. S. S. J. Rewell, L. Churilov, T. K. Sidon, E. Aleksoska, S. F. Cox, M. R. Macleod, D. W. Howells, Evolution of ischemic damage and behavioural deficit over 6 months after MCAo in the rat: Selecting the optimal outcomes and statistical power for multi-centre preclinical trials. PloS One 12, e0171688 (2017).

43. M. J. Haley, C. B. Lawrence, The blood–brain barrier after stroke: structural studies and the role of transcytotic vesicles. J. Cereb. Blood Flow Metab. 37, 456–470 (2017).

44. V. S. Polikov, P. A. Tresco, W. M. Reichert, Response of brain tissue to chronically implanted neural electrodes. J. Neurosci. Methods 148, 1–18 (2005).

45. J. C. Chen, P. Kan, Z. Yu, F. Alrashdan, R. Garcia, A. Singer, C. S. E. Lai, B. Avants, S. Crosby, Z. Li, B. Wang, M. M. Felicella, A. Robledo, A. V. Peterchev, S. M. Goetz, J. D. Hartgerink, S. A. Sheth, K. Yang, J. T. Robinson, A wireless millimetric magnetoelectric implant for the endovascular stimulation of peripheral nerves. *Nat*. Biomed. Eng. 6, 706–716 (2022).

46. E. T. Zhao, J. Hull, N. M. Hemed, H. Uluşan, J. Bartram, A. Zhang, P. Wang, A. Pham, S. Ronchi, J. R. Huguenard, A. Hierlemann, N. A. Melosh, A CMOS-based highly scalable flexible neural electrode interface. bioRxiv, 2022.11.03.514455 (2022).

47. Y. Kim, G. A. Parada, S. Liu, X. Zhao, Ferromagnetic soft continuum robots. Sci. Robot. 4, eaax7329 (2019).

48. J. M. Lee, G. Hong, D. Lin, T. G. Schuhmann Jr, A. T. Sullivan, R. D. Viveros, H.-G. Park, C. M. Lieber, Nanoenabled direct contact interfacing of syringe-injectable mesh electronics. Nano Lett. 19, 5818–5826 (2019).

