## Supplementary Information for "Ultra-flexible endovascular probes for brain recording through micron-scale vasculature"

This file contains:

Supplementary text

Supplementary Figures S1 – S13

**Bending Stiffness of Micro-Endovascular (MEV) Probes**

To calculate the bending stiffness of the MEV probe, here we only consider the guidewire made of a 10 μm thick SU8 layer and not the 0.8 μm thick mesh-like structure, which supports the recording electrodes, on either side of the guide (Fig. 1). Specifically, the cubic proportionality of bending stiffness to thickness implies that the guide will be >1000 times stiffer than surrounding mesh structure. To model the deflection, we use a beam of length $L$ = 18 mm, the length of the device region, thickness $t$ = 10 μm, width $b$ = 25 or 75 μm. The face of the beam that is on the yz-plane and at x = 0 is fixed, and the yz-plane face that is located at x = $L$ has a force $F$ applied along the z-direction.

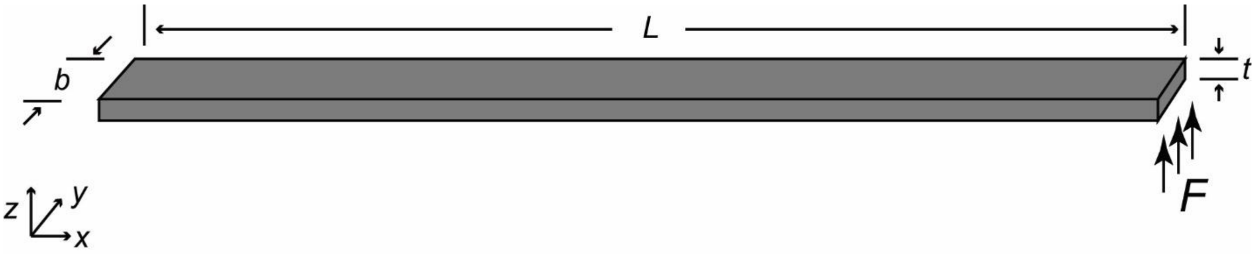


The area moment of inertia, $I$, can be estimated as

$$I = \frac{t^3 b}{12}$$

The deformation, $w$, at x = $L$ can be expressed as

$$w = \frac{4FL^3}{Et^3 b}$$

where $E$ = 2 GPa is Young's modulus of SU-8.

The bending stiffness, $k$, can be expressed as

$$k = \frac{F}{w} = \frac{Et^3 b}{4L^3}$$

1) For the 25-μm guidewire in the device region, $b$ = 25 μm, which yields $k = 2.14\times10^{-6}$ N/m.
2) For the 75-μm guidewire in the device region, $b$ = 75 μm, which yields $k = 6.43\times10^{-6}$ N/m.

Therefore, under the same applied force $F$, deflection of the tip of the 25-μm guidewire probe is 3 times of that of the 75-μm guidewire probe, consistent with the expected linear scaling in beam width. In comparison, the SU-8 ribbons in the mesh structure are 800 nm in thickness and 10 μm in width, which yields $k = 4.39 \times10^{-10}$ N/m, and confirmed that the guidewire is >1000 times stiffer than surrounding mesh structure.

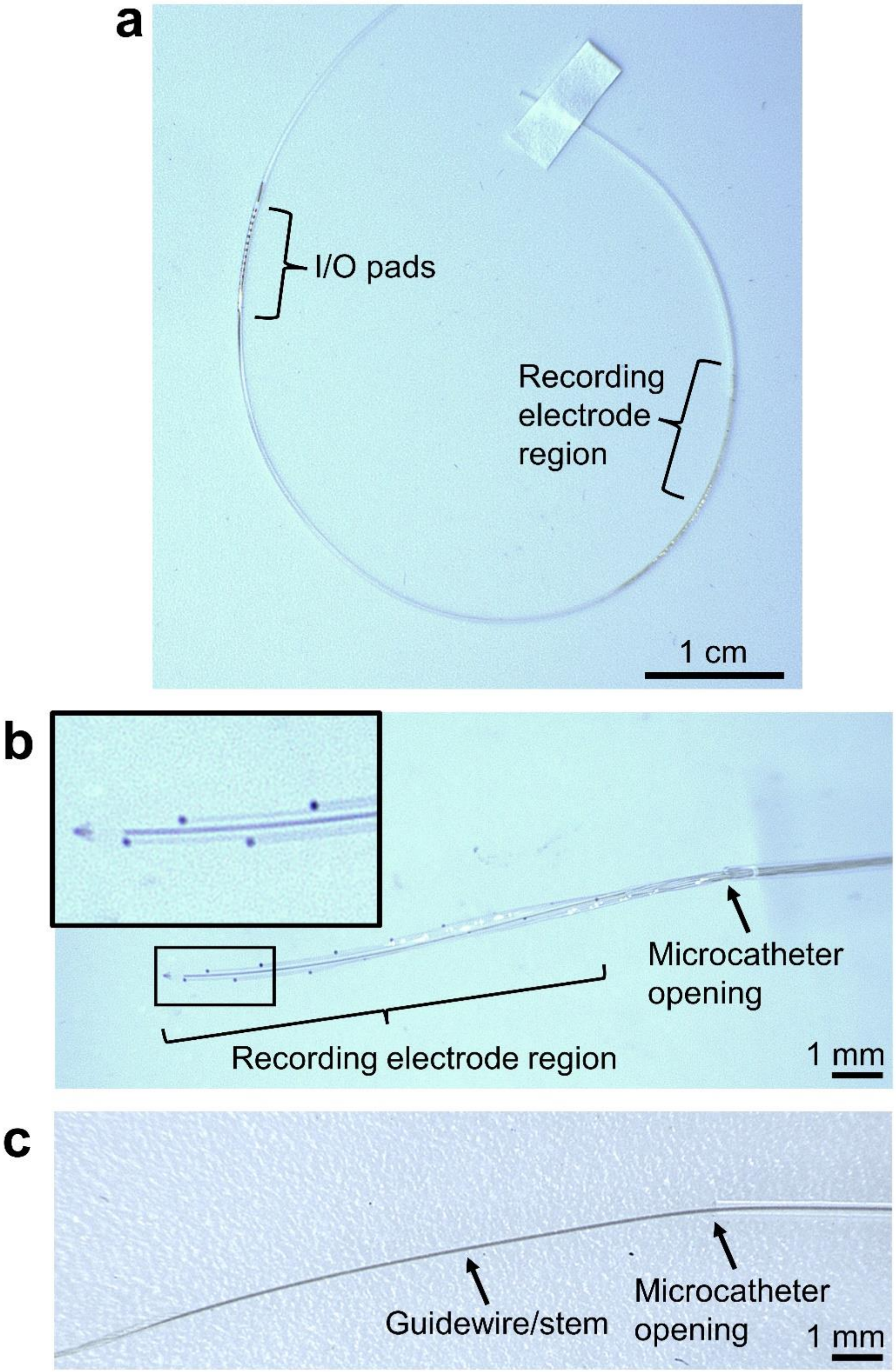


**Supplementary Figure S1 | Loading and injection of MEV probes. a**, A probe with a 75-µm wide guidewire loaded into the flexible microcatheter with the inner diameter of 200 µm and the outer diameter of 350 µm. **b**, The 900-µm-wide device region relaxes and unfolds in water after injection from the microcatheter. Inset shows the electrodes at the probe tip. **c**, The stem region coated with the guidewire is subsequently injected from the microcatheter.

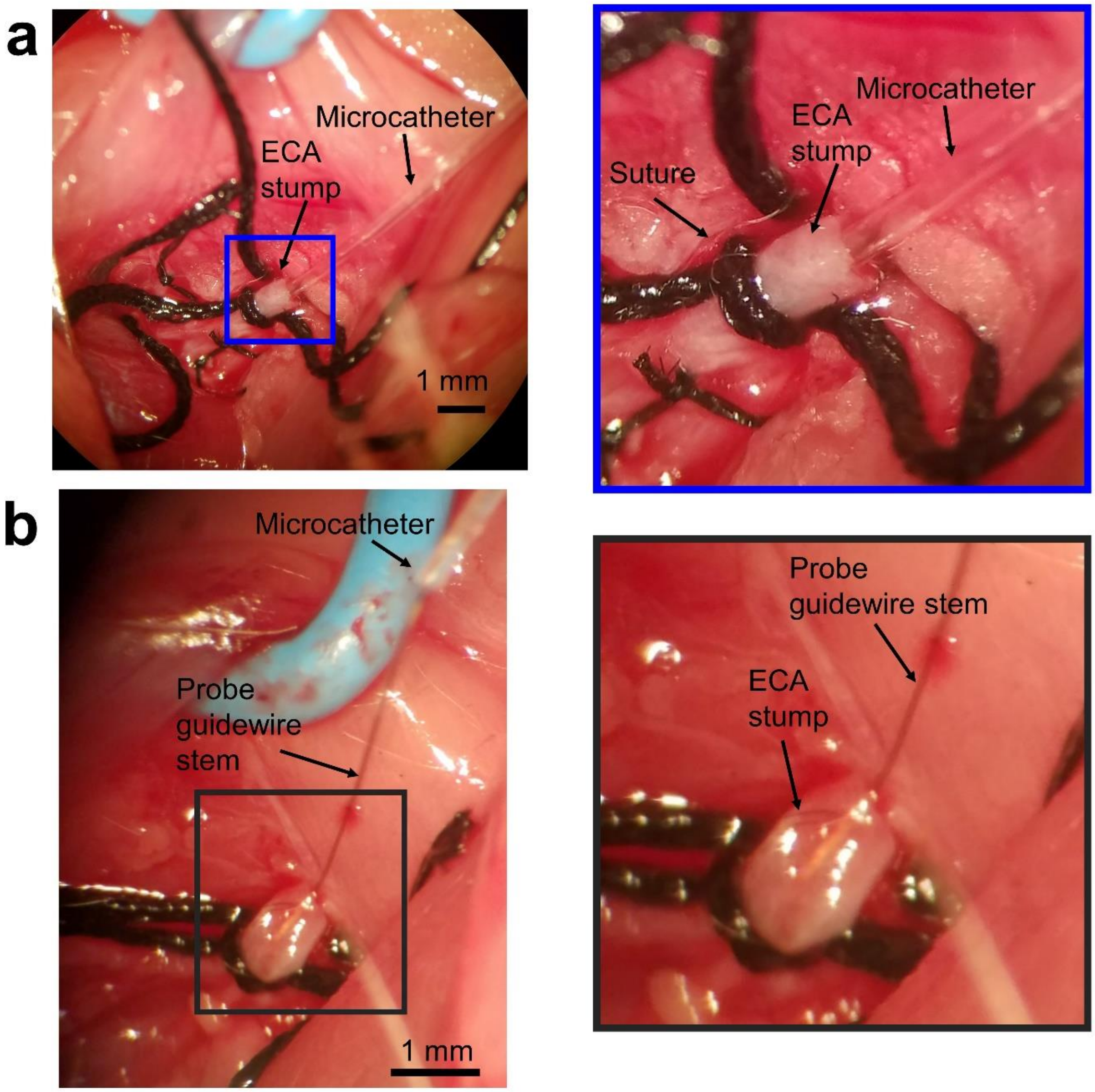


**Supplementary Figure S2 | Implantation procedure. a**, Microcatheter loaded with an MEV probe (not shown) is inserted into ECA stump and advanced to the MCA/ACA bifurcation. A suture is placed around the ECA stump to prevent blood leakage around the microcatheter. **b**, After injection, the microcatheter is retracted, the guidewire stem is exposed and fixed by tying the suture around the ECA stump.

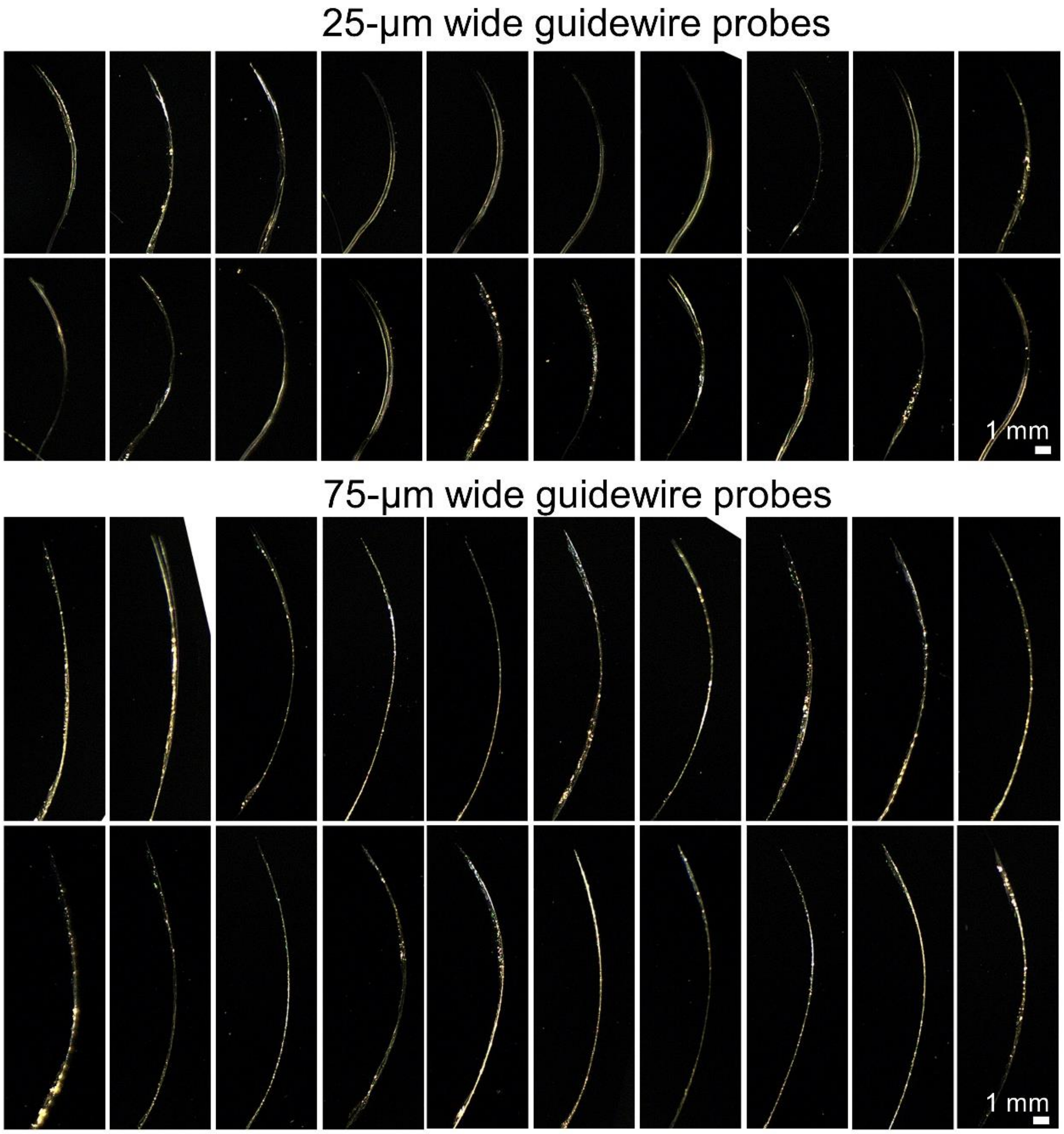


**Supplementary Figure S3 |** Side view of the bending of the 18-mm long device region of the MEV probes (N = 20 for 25-µm and 75-µm wide guidewires, guidewire thicknesses are consistently 10 µm) in saline due to the residue stress of SU-8 after release of the probes from the fabrication substrate. These images are used to calculate the radii of curvature and effective bending angles in Fig. 2c. Probes with 25-µm and 75-µm wide guidewires can be selectively implanted into the MCAs and the ACAs, respectively, as the MCA and the ACA branches form different angles with the ICA.

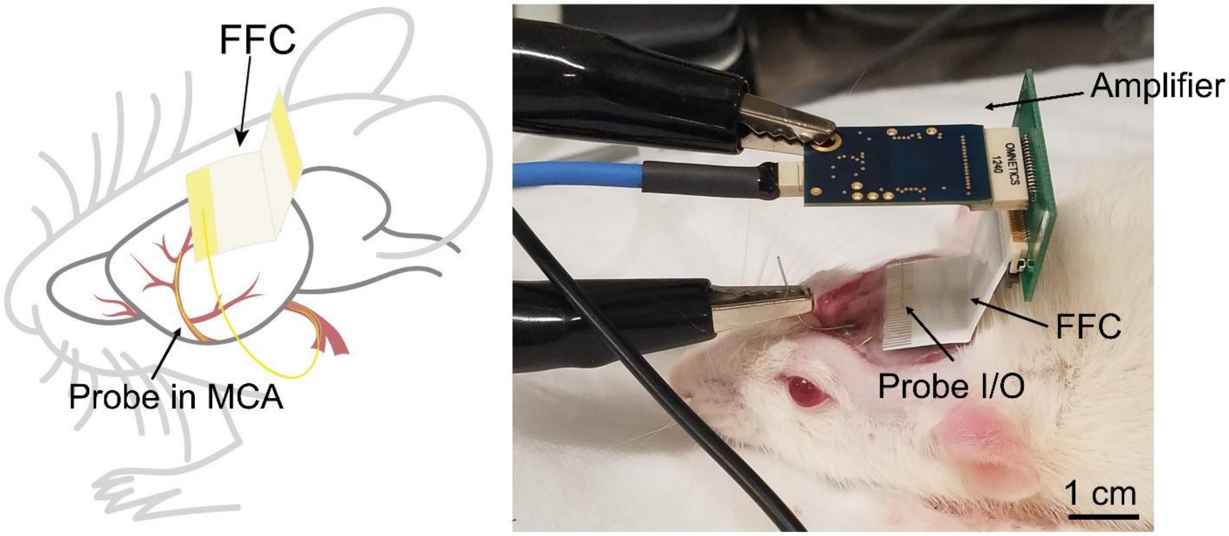


**Supplementary Figure S4 | Electrophysiology recording setup.** Schematic (left) and photo (right) showing that an MEV probe implanted into a rat brain is threaded through a subcutaneous tunnel underneath the skin around the head, and the I/O region is aligned onto the conductors of the FFC fixed on the skull. The FFC is connected to an amplifier and an Intan RHD 2132 amplifier evaluation system connected to a recording computer (not shown).

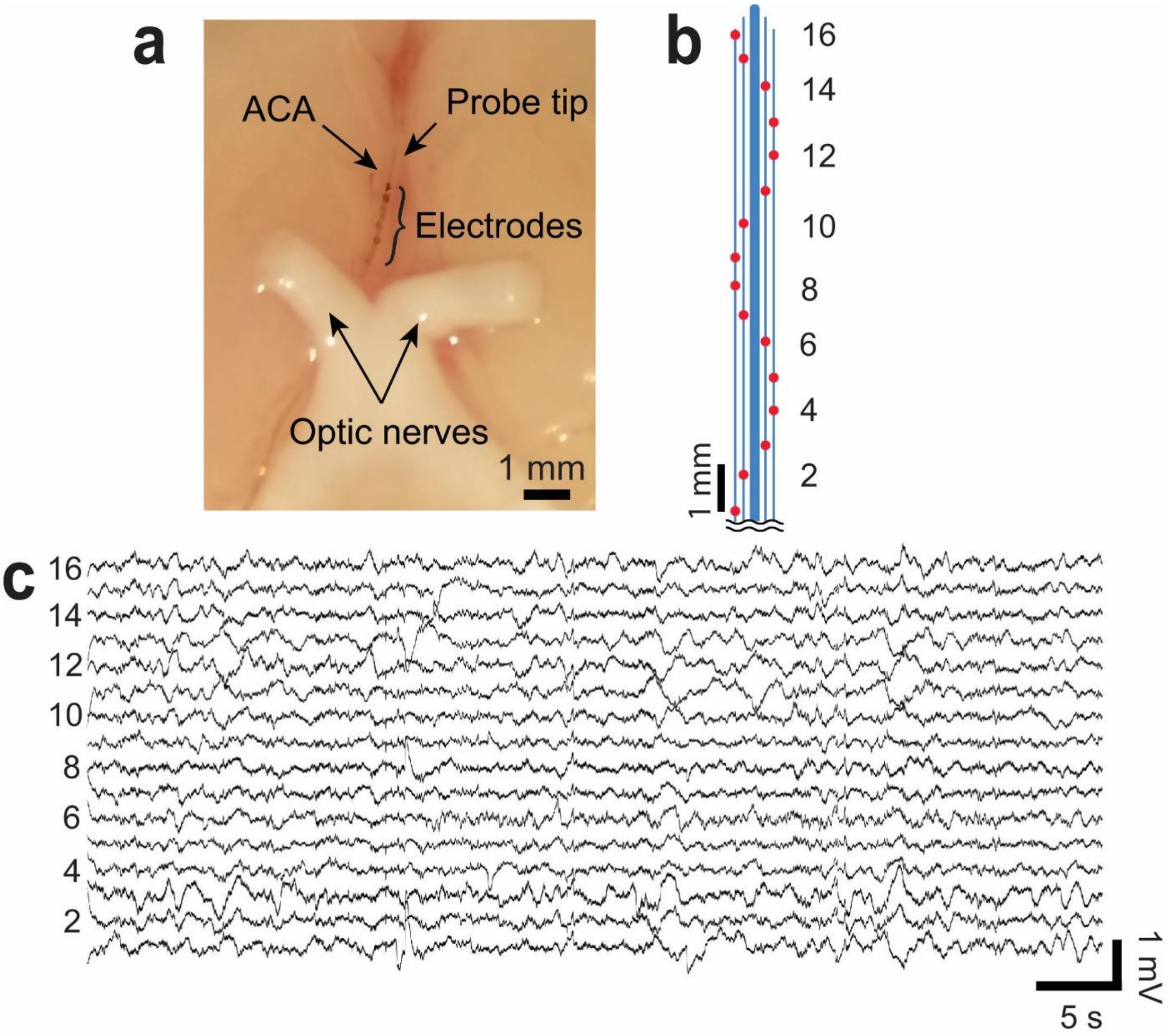


**Supplementary Figure S5 | In vivo endovascular recording in ACA. a**, Rat brain with an MEV probe in the ACA. (**b**) Schematic of the probe device region showing the relative positions of 16 Pt electrodes marked by red spots, higher-numbered channels were implanted deeper in the ACA. (**c**) The corresponding acute in vivo 16-channel unfiltered recording showing local field potential oscillations.

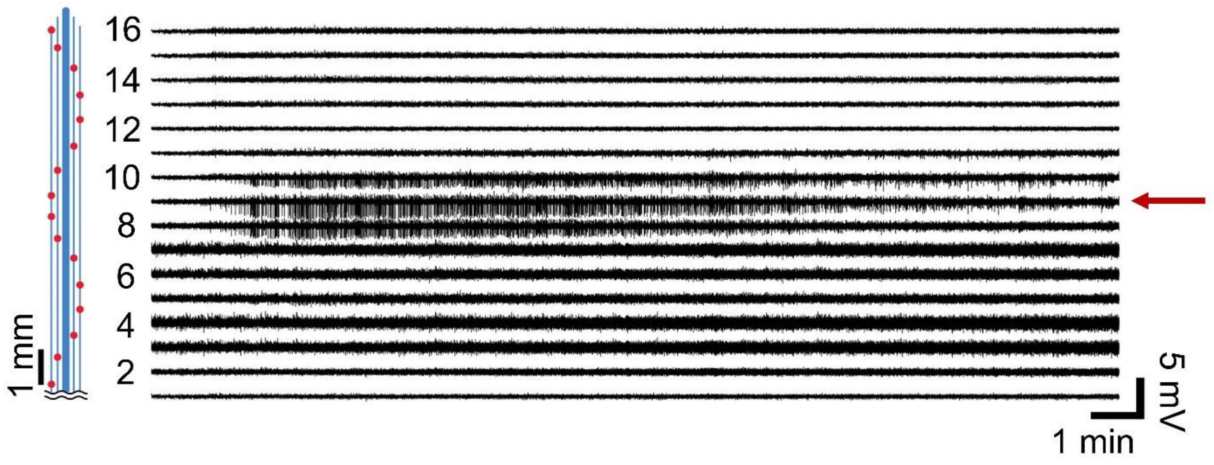


**Supplementary Figure S6 | Seizure recording in all 16 channels from the MEV probe in the MCA.** Data shown is the unfiltered recording showing seizure activity from the MEV probe in Fig. 3c. The relative positions of 16 Pt electrodes are marked by red spots in the schematic on left. Seizure spikes were recorded by three adjacent channels (Ch. #8 – Ch.#10), demonstrating the ability of the MEV probes to locate and track the seizure foci. The channel plotted in Fig. 3c is highlighted by the red arrow.

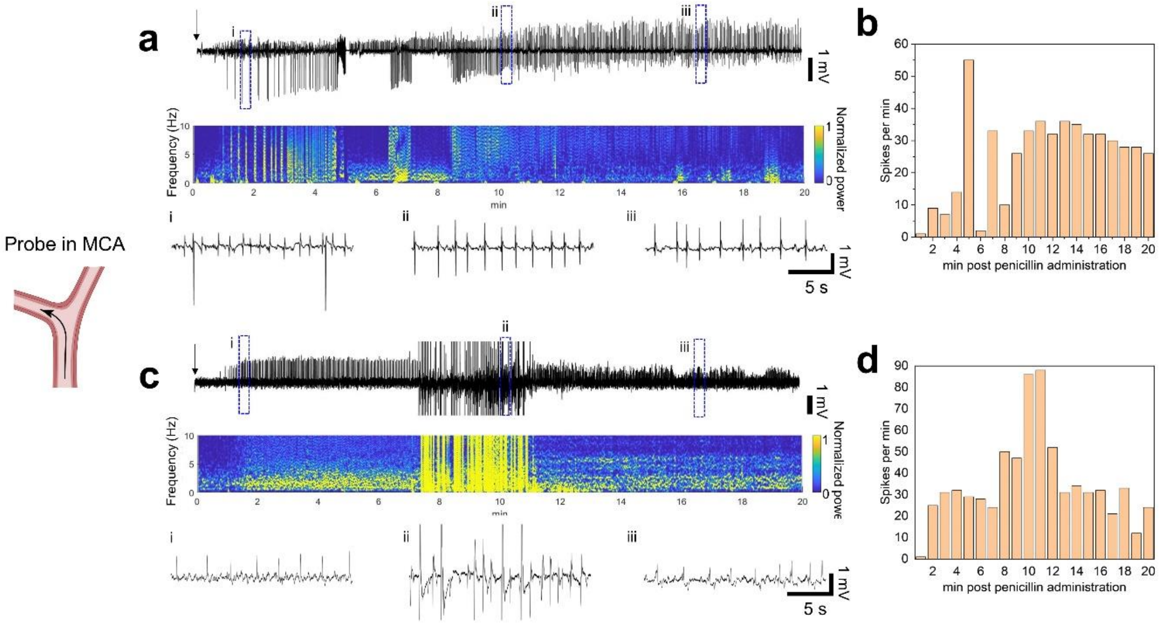


**Supplementary Figure S7 | Seizure recording from two additional rats with MCA probes.** **a**,**c**, Top, penicillin-induced seizures recorded by a representative channel from probes in MCA for 20 min. Middle, spectrogram post penicillin injection. The colormap shows the normalized power levels. Bottom, zoom-in views showing the evolution of seizure spikes at three different time points highlighting the changes in spike amplitude over time. Black arrows denote the time point of penicillin injection. **b**,**d**, Number of seizure spikes per min recorded by probes in MCA. The probe in (**b**) recorded 505 spikes, and the probe in (**d**) recorded 711 spikes in 20 min.

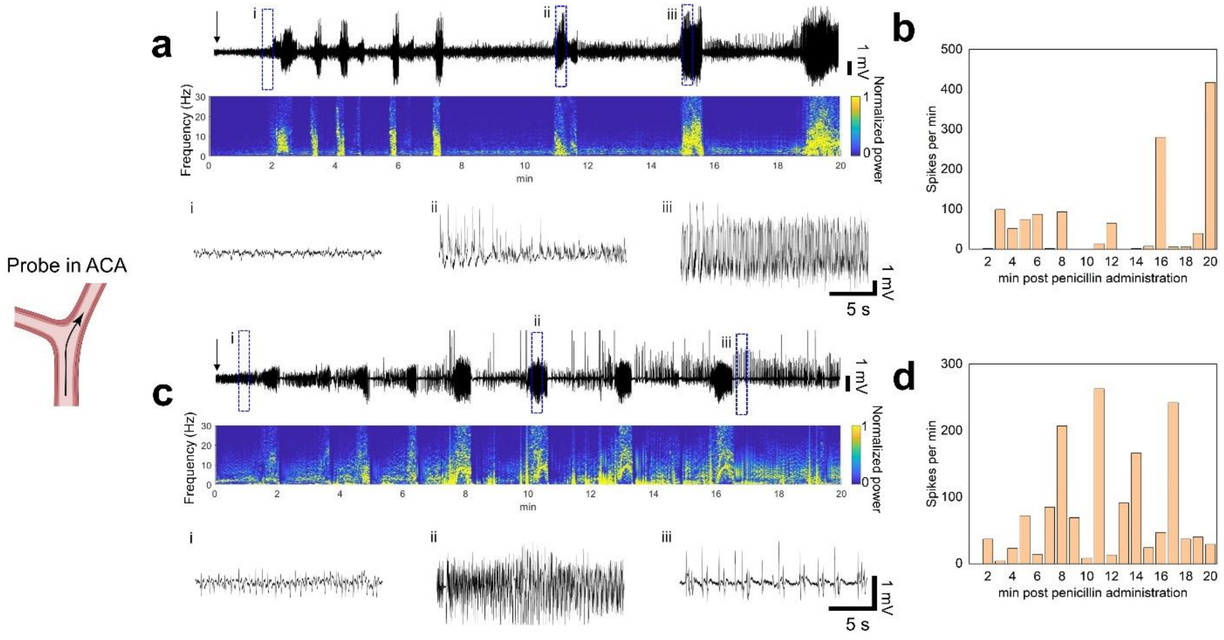


**Supplementary Figure S8 | Seizure recording from two additional rats with ACA probes.** **a**,**c**, Top, penicillin-induced seizures recorded by a representative channel from probes in ACA for 20 min. Middle, spectrogram post penicillin injection. The colormap shows the normalized power levels. Bottom, zoom-in views showing the evolution of seizure spikes at three different time points highlighting before and during the burst firing activity. Black arrows denote the time point of penicillin injection. **b**,**d**, Number of seizure spikes per min recorded by probes in ACA. The probe in (**b**) recorded 1238 spikes, and the probe in (**d**) recorded 1471 spikes in 20 min.

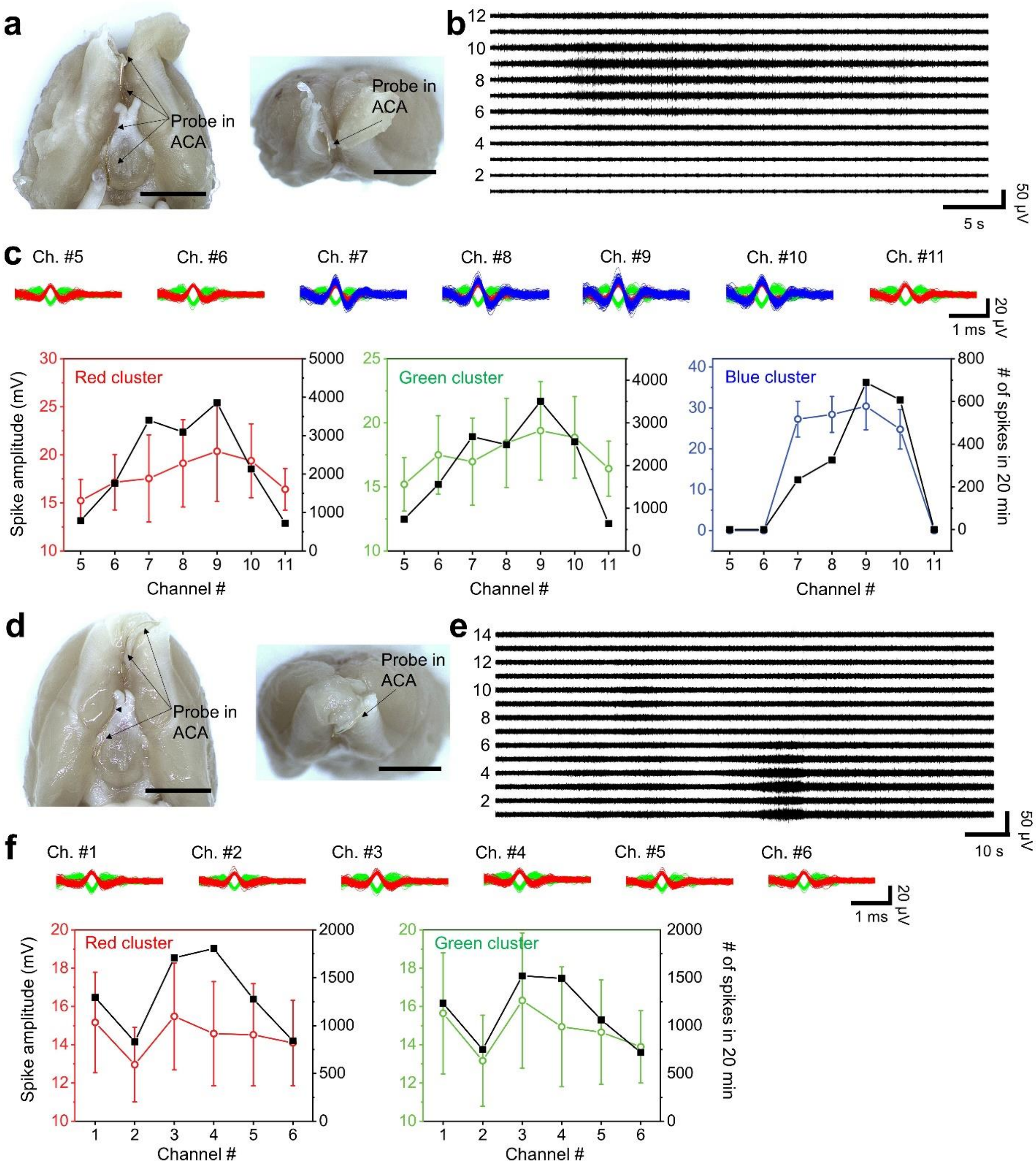


**Supplementary Figure S9 | Additional single unit recording data. a-b**, Bottom (left) and front (right) views of a rat brain with an MEV probe in ACA in the olfactory bulb (**a**) and the corresponding multi-channel recording from the same probe after 250–6,000 Hz band-pass filtering to show single unit burst activity (**b**). Higher-number channels were implanted deeper in the ACA. Scale bars in **a** are 5 mm. **c**, Top, sorted spikes assigned to different neurons from the channels with single unit activity. Each distinct color in the sorted spikes represents a unique identified neuron. Bottom, spike amplitude (with ±1 standard deviation, s.d.) and the number of single unit spikes recorded in 20 min. All neurons exhibit higher spike amplitude and number of spikes in Ch. #9 and decaying in the channels on either side of the probe, indicating that Ch. #9 is the closest to the spiking neurons. **d-f**, Similarly recorded and analyzed data from another rat. Neurons exhibit higher spike amplitude and number of spikes in Ch. #3 and #4 and decaying in the channels on either side of the probe, indicating that Ch. #3 and #4 are the closest to the spiking neurons.

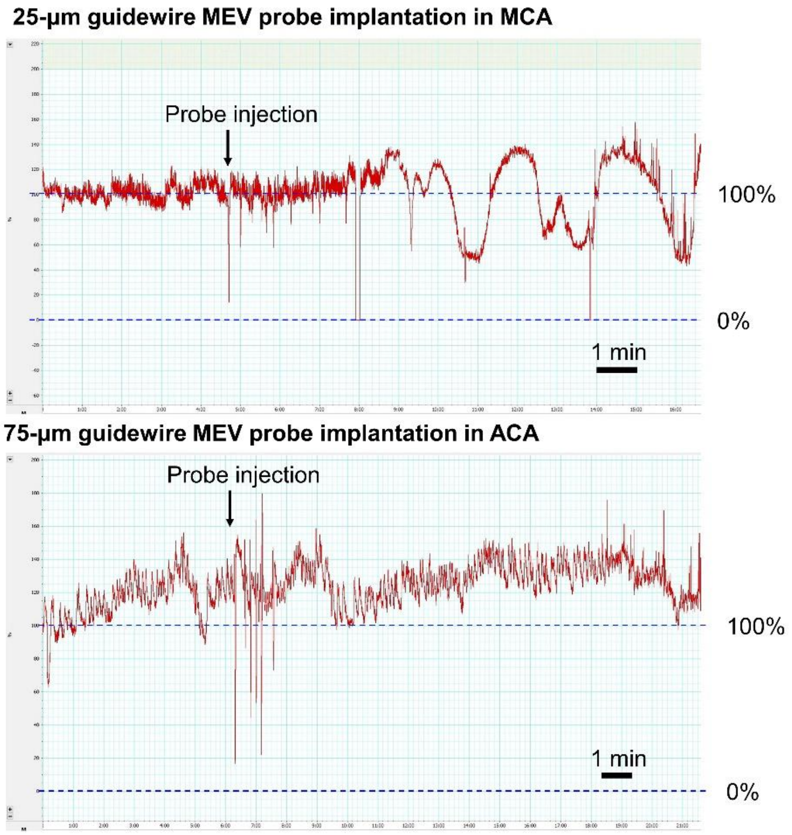


**Supplementary Figure S10 | Representative laser doppler flowmetry of probe implantation in MCA and ACA.** After injection of a probe with 25-μm wide guidewire in MCA, the cerebral blood flow fluctuated between 60% to 140%, while after injection of a probe with 75-μm wide guidewire in ACA, the cerebral blood flow did not change. The probes have the same support structure except for the guidewires. The injection depth exceeds 1 cm past the MCA/ACA bifurcation. The post-implantation blood flow levels were normal, and the rats did not suffer stroke or any other neurologic deficit from the surgery.

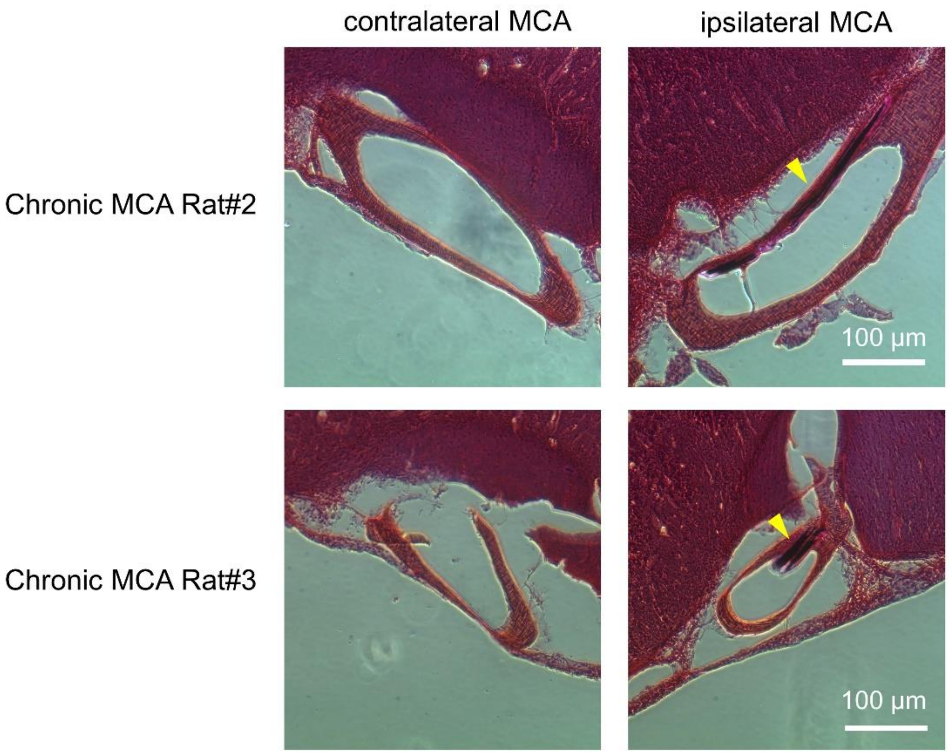


**Supplementary Figure S11 |** MCA cross-sections from the H&E-stained slices from two additional rats 28 days post-implantation in MCA. The yellow arrow highlights the position of the MEV probes in the vessel wall.

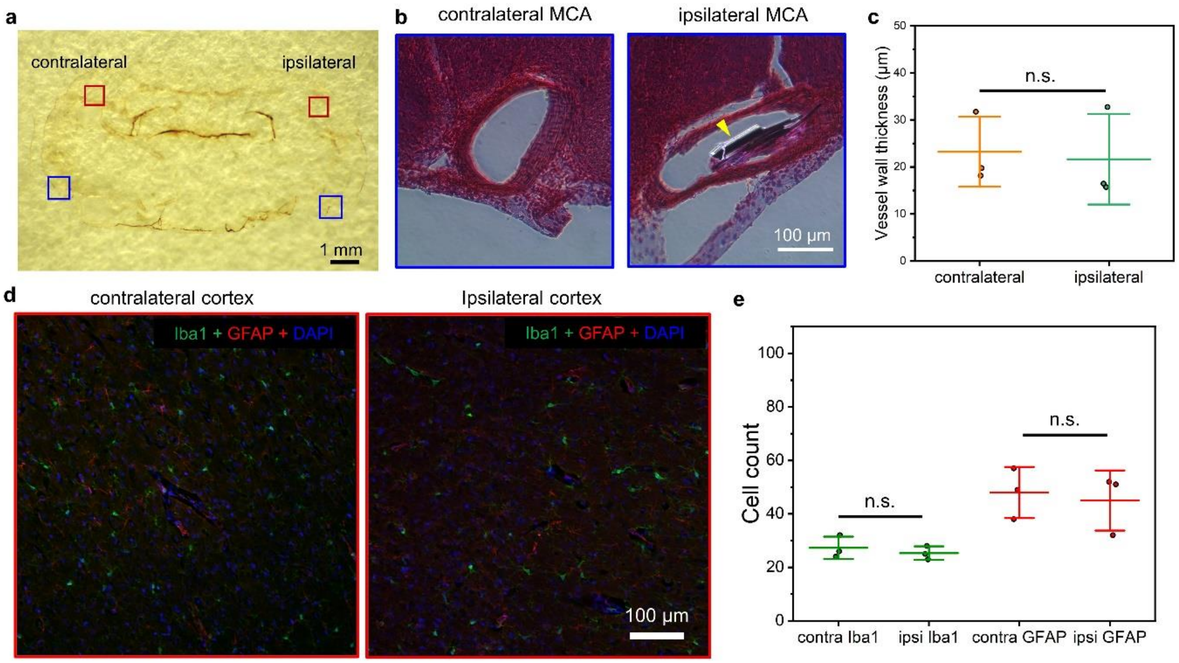


**Supplementary Figure S12 | Short-term histology 3 days post implantation of an MEV probe in MCA. a**, Digital camera image of a representative IgG-stained brain slice 3 days post-implantation in MCA. **b**, Zoom-in views of the contralateral and ipsilateral MCA cross-sections from the H&E-stained slice from the regions highlighted by the blue boxes in **a**. The yellow arrow highlights the probe in the vessel. **c**. MCA vessel wall thickness measured from H&E images of 3 brain slices that are 600 μm apart (with ±1 standard deviation, s.d.). **d**, Confocal fluorescence microscopy images of the contralateral and ipsilateral cortexes from the regions highlighted by the red boxes in **a**. The brain slice was stained with antibodies for Iba1 (green) and GFAP (red), and DAPI (blue). **e**, Number of microglia (Iba1) and astrocytes (GFAP) counted from fluorescence images of 600 μm*450 μm from 3 brain slices that are 600 μm apart (with ±1 standard deviation, s.d.). n.s.= nonsignificant; unpaired two-tailed t test.

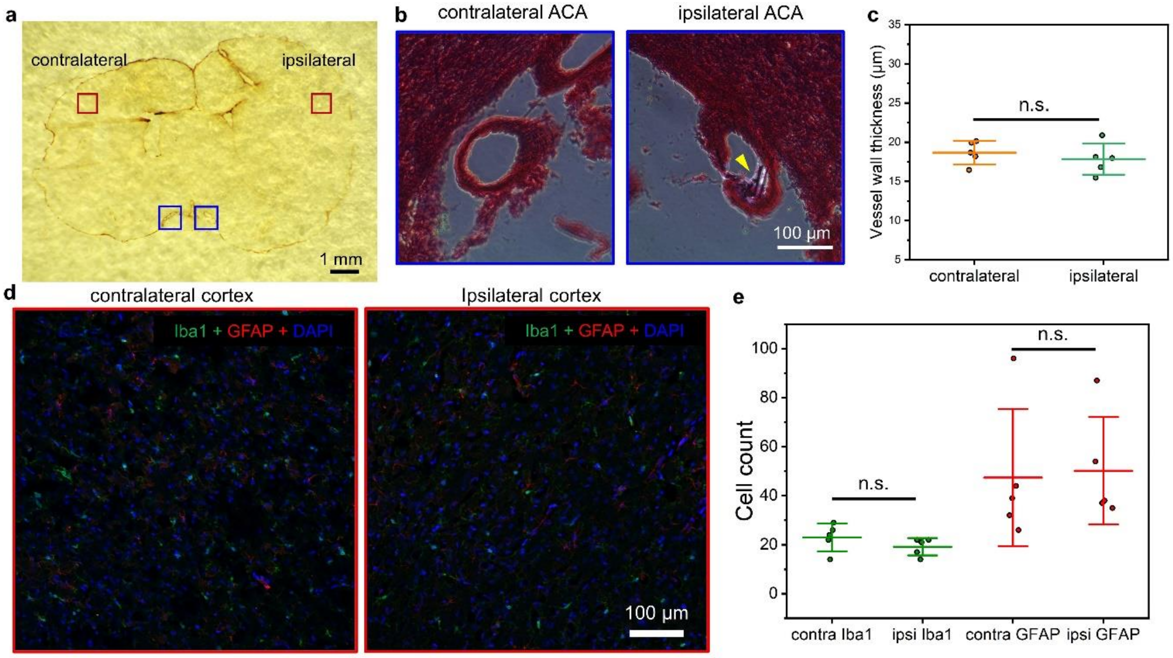


**Supplementary Figure S13 | Short-term histology 3 days post implantation of an MEV probe in ACA. a**, Digital camera image of a representative IgG-stained brain slice 3 days post-implantation in ACA. **b**, Zoom-in views of the contralateral and ipsilateral ACA cross-sections from the H&E-stained slice from the regions highlighted by the blue boxes in **a**. The yellow arrow highlights the probe in the vessel. **c**. ACA vessel wall thickness measured from H&E images of 3 brain slices that are 600 μm apart (with ±1 standard deviation, s.d.). **d**, Confocal fluorescence microscopy images of the contralateral and ipsilateral cortexes from the regions highlighted by the red boxes in **a**. The brain slice was stained with antibodies for Iba1 (green) and GFAP (red), and DAPI (blue). **e**, Number of microglia (Iba1) and astrocytes (GFAP) counted from fluorescence images of 600 μm*450 μm from 3 brain slices that are 600 μm apart (with ±1 standard deviation, s.d.). n.s.= nonsignificant; unpaired two-tailed t test.
